# Dissecting the Complexities of Learning With Infinite Hidden Markov Models

**DOI:** 10.1101/2023.12.22.573001

**Authors:** Sebastian A. Bruijns, International Brain Laboratory, Kcénia Bougrova, Inês C. Laranjeira, Petrina Y. P. Lau, Guido T. Meijer, Nathaniel J. Miska, Jean-Paul Noel, Alejandro Pan-Vazquez, Noam Roth, Karolina Z. Socha, Anne E. Urai, Peter Dayan

## Abstract

Learning to exploit the contingencies of a complex experiment is not an easy task for animals. Individuals learn in an idiosyncratic manner, revising their approaches multiple times as they are shaped, or shape themselves, and potentially end up with different strategies. Their long-run learning curves are therefore a tantalizing target for the sort of individualized quantitative characterizations that sophisticated modelling can provide. However, any such model requires a flexible and extensible structure which can capture radically new behaviours as well as slow changes in existing ones. To this end, we suggest a dynamic input-output infinite hidden semi-Markov model, whose latent states are associated with specific components of behaviour. This model includes an infinite number of potential states and so has the capacity to describe substantially new behaviours by unearthing extra states; while dynamics in the model allow it to capture more modest adaptations to existing behaviours. We individually fit the model to data collected from more than 100 mice as they learned a contrast detection task over tens of sessions and around fifteen thousand trials each. Despite large individual differences, we found that most animals progressed through three major stages of learning, the transitions between which were marked by distinct additions to task understanding. We furthermore showed that marked changes in behaviour are much more likely to occur at the very beginning of sessions, i.e. after a period of rest, and that response biases in earlier stages are not predictive of biases later on in this task.

## 1 Introduction

Engaging with a new environment or experiment raises a multitude of questions: which sensory signals are pertinent to the task at hand, and which are just noise? What actions are relevant to performance? How should observations inform actions? How can reward be maximized? Particularly if the experimenter changes the task suddenly in order to shape behaviour, but also in stable environments, animals solve these problems through a mixture of apparent leaps in performance and slow accumulation of improvements (Breland & Breland, 1951; Epstein, Kirshnit, Lanza, & Rubin, 1984; Gallistel, Fairhurst, & Balsam, 2004; Köhler, 1948; Krueger & Dayan, 2009; Luft & Buitrago, 2005; Maier, 1931; Moore & Kuchibhotla, 2022; Rescorla, 1972). This process of learning is marked by substantial variability across individuals, who progress at different speeds and over distinct intermediate stages (Piaget, 1952). Even if the ultimate behaviour is indistinguishable, this initial variability can make comparisons across groups during learning challenging (The International Brain Laboratory et al., 2021). More generally, the particularities of the learning path may never be fully overcome, so that their trace is detectable in performance even after learning has finished and behaviour has stabilised (Dayan, Roiser, & Viding, 2020).

Despite the richness and importance of these dynamics, much of the work on the modelling of learning has ignored this sort of acquisition, generally considering the adjustments that occur to adapt to ongoing changes in modest facets of tasks, such as reversing reward schedules. By this point, most of the problem has been solved, or subjects who failed to learn have been discarded. One of the main reasons for this is that each animal provides only one sample of a learning curve, whereas for fully acquired behaviour, every trial can typically be viewed as another sample from the learned behaviour. This means that learning curve data are generally sparse, further aggravating the problem of large variability. Here, we make use of the large-scale approach to data collection embodied by the International Brain Laboratory (The International Brain Laboratory et al., 2021), and base our analysis on the multi-session learning curves of more than 100 mice coming to solve a perceptual decision-making task.

Nevertheless, there has been some work on quantifying acquisition. One set of results concerns the point in time at which an animal can be said to have “learned” a task, often defined as reliably above chance performance (Smith et al., 2004). This kind of change-point detection is a difficult challenge, as behaviour is generally probabilistic and can be erratic, prompting the development of a number of approaches (see e.g. Jang et al. (2015); Papachristos and Gallistel (2006)). These methods are however more concerned with finding changes in general, in particular making a binary distinction between uninformed and learned behaviour, rather then describing used strategies in detail, or finding possible intermediate stages. Recent work addressing strategy inference more specifically, which does consider learning, includes the approach of Maggi et al. (2022). They combined a set of simple, pre-selected strategies with an inference mechanism for performing strategy inference on a trial-by-trial basis. By exponentially decaying evidence over time, they were able to track the arrival and departure of various strategies. Their framework does however require formalising possible strategies ahead of time, and cannot incorporate probabilistic strategies. There has also work on forms of progressive meta-learning in humans (Jain et al., 2023).

To accommodate the complexities of learning curves, a descriptive modelling framework must fulfil a number of desiderata. First, at any point along the curve, it should capture the current repertoire of behaviours, characterizing performance. Second, it needs to track this repertoire as behaviour evolves, introducing new components (which we identify as behavioural ‘states’) when change is abrupt (e.g. Durstewitz, Vittoz, Floresco, and Seamans (2010); Gallistel et al. (2004)), detecting the re-use of a past state if it re-emerges, and allowing for slow, gradual shifts in a component, for instance, with the steady development of skilled performance (e.g. Song, Baah, Cai, and Niv (2022)). A third requirement is that the collection of components should be potentially unbounded, since it is not generally possible to pin down ahead of time how many distinct behaviours any individual animal might exhibit. Finally, it should be possible to switch between the states within a current repertoire, as rapidly as from one trial to the next.

We satisfy all three requirements by building a model that combines and extends two recent approaches. One is from Ashwood et al. (2022) (see also Calhoun, Pillow, and Murthy (2019)), who described decision-making performance after learning with a hidden Markov model (HMM). In this, each hidden or latent state of the model captures a single component of behaviour in the form of a map from task-relevant variables to distributions over choices, for instance via logistic regression. In the case of perceptual decision-making, this generalizes a psychometric function to include factors such as perseveration, by considering previous choices as an input feature. The overall description of behaviour is in terms of a mixture of different policies (such as engaged or disengaged motivational states) that can switch rapidly. However, the HMM approach of Ashwood et al. (2022) assumes stationarity of behaviour across time, and is constrained to a fixed level of complexity, by specifying the number of states, and thereby the number of parameters, *a priori*. This makes it ill-suited to characterising the dynamic and idiosyncratic progression through training. To address these issues, we adopted the HMM framework to capture abrupt changes, except that (i) along with the motivational factors, the latent states can describe what the animal knows about the task at any point; that (ii) we used a semi-Markov model so that latent states can persist for non-exponentially distributed numbers of trials; and that (iii) the states come from a Bayesian non-parametric structure, allowing for a degree of behavioural complexity that is only constrained by an inbuilt Occam’s razor, and enabling the introduction of new states for suddenly appearing new behaviours (Beal, Ghahramani, & Rasmussen, 2001; Gershman & Blei, 2012; Heald, Lengyel, & Wolpert, 2021; Johnson & Willsky, 2013; Teh, Jordan, Beal, & Blei, 2006).

The second approach is that of Roy, Bak, Akrami, Brody, and Pillow (2021), which effectively considers just a single state, but allows the weights of the logistic regression to be dynamic, tracking changes in behaviour through appropriate changes in the weights. We used this so that the characteristics of our hidden states can evolve slowly, capturing the other prevalent form of acquisition of skilled performance.

Using our combined model, we show that learning progressed over a small number of distinct stages which are present in almost all animals. These stages apparently correspond to the sequential acquisition of elements of the task – in our case, particularly associated with taking into account different aspects of the sensory environment inherent to the task. Although this pattern was shared across all the mice, the duration and diversity of the stages differed greatly between individuals. We also found other prevalent patterns, including regression from newer states affording good performance to older ones affording worse performance over the course of the learning curve.

We first describe the IBL task and our way of characterising the behaviour mice exhibit; then discuss the details of the model by studying a representative fit to one animal in detail; and conclude by summarising the fits of our model to 119 subjects, highlighting similarities and differences across the population.

## 2 Results

We analysed the choices of 119 mice learning a perceptual decision-making task, each of them going through on average 24.2 (total: *>*2800) sessions and on average *∼*14700 trials (total: *>*1.7 million) (The International Brain Laboratory et al., 2021). In this task, head-fixed mice were shown a sinusoidal grating of a controlled contrast, with equal probability on either the right or left side of a screen. They then had to center it (within 60 s) by turning a steering wheel in the appropriate direction. Successful trials led to water reward; unsuccessful trials to a noise burst and a 1 s timeout. Trials were self-paced, with mice signalling their readiness by keeping the wheel still for a period.

Mice learned the task according to a rigorous shaping protocol, which introduced more difficult stimuli gradually and actively removed action biases. Accordingly, shaping started with gratings of the highest contrasts: 100% and 50%. At this initial stage, there was no perceptual difficulty, but the animals had to learn the basic contingencies and requirements of the task. Once they had reached sufficient performance on these contrasts (≥80% correct for each contrast type on the last 50 trials), 25% strength contrasts were introduced. After performance was good on this extended set as well (same criterion for each contrast), the remaining contrasts were introduced in a staggered manner: 12.5%, 6.125%, and 0%, while the 50% contrast was dropped from the task. For the 0% contrast, one side was rewarded randomly (with probabilities 50% each). A debiasing protocol increased the probability of repeating the stimulus that was just shown when the mouse made a mistake on an easy (100% or 50%) contrast. This served to motivate the animals away from perseverative or biased strategies, and could lead to reward rates lower than 50% (as would otherwise be expected from pure chance).

In order to characterise the course of learning across every trial, we developed a flexible model which segments the behaviour of an animal into discrete states that last for variable numbers of trials within a session and can recur in multiple sessions. As this is a descriptive model, we equate a behaviour with its corresponding state, and generally will not distinguish between the two in the text. We first describe how a single state generates choice probabilities on a trial for which it was responsible (**Fig. 1** within circles); and then how we treat multiple states (**Fig. 1** arrows).

**Fig. 1.**
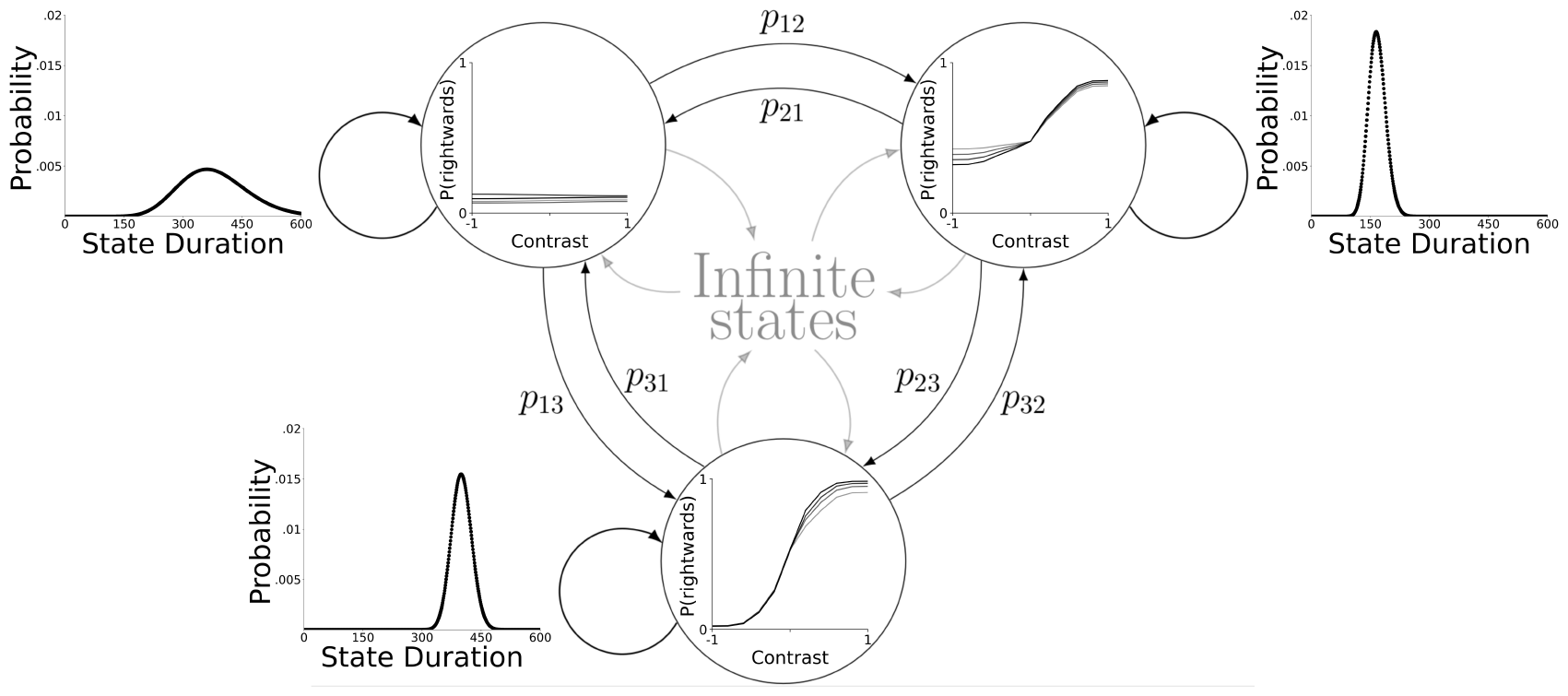
Visual representation of the main components of the model. Each state, represented by a circle, has an observation distribution associated with it, shown inside its circle. This distribution is implemented via logistic regression, to consider the contrast of the current trial and a weighted history of previous choices (the latter is not shown here). The weights underlying these regressions can change from session to session, resulting in shifts of the psychometric functions (PMFs) they represent; we depict this evolution here with varying shades of grey. States are connected to other existing states via transition probabilities *p*. In addition to that, states also have the option to transition into a new state, to describe a type of behaviour which is not well captured by any of the existing states. Lastly, staying in the same state for more than one trial is not modelled via a self-transition probability, but instead each state has its own duration distribution, which determines for how many trials it remains active.

As in previous work (Ashwood et al., 2022; Roy et al., 2021), we formalise the response probabilities for the binary choices of mice through logistic regression (omitting the rare trials in which the animal timed out by not responding within 60 s). Trial *t* of session *n* is described by features ***f***_*n,t*_ comprising: (i) the stimulus, i.e. the contrast on the left and right of the screen; separated, to allow for different sensitivities to leftwards and rightwards stimuli, which is important because mice were frequently differently sensitive to the screen sides in this task (ii) task history, in the shape of an exponentially decaying average of the last 5 actions; since mice appear not to have used reward information to implement win-stay lose-shift strategies, but rather employed a perseverative bias to repeat previous actions (as was also observed in Miller, Botvinick, and Brody (2021), Beron, Neufeld, Linderman, and Sabatini (2022), and, in the same task at a later stage, by Findling et al. (2023)) (iii) a bias term to allow for side preferences regardless of other features. If we label the state that is active on this trial as *x*_*n,t*_, the response *y*_*n,t*_ *∈ {*L, R} (for Left and Right) is modelled by the distribution

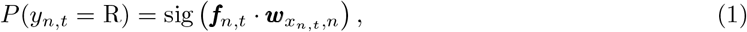

where the weights of the states ***w***_*x,n*_, ∀*x* are also indexed by session *n* as they can drift across sessions. Here sig(*·*) is the standard logistic sigmoid function.

The model generalises a standard hidden Markov model (HMM) in three ways that make it especially suited to describe the phases of learning, see also **Fig. 1**: (i) it is non-parametric about the number of states, i.e. the number of states appropriate for describing the behaviour of each individual is separately determined, thus accommodating inter-individual differences. This characteristic also allows the model to capture sudden changes in behaviour, as it is able to introduce a new state when behaviour changes systematically (we call this the ‘fast process’). (ii) States are dynamic over sessions *n* (see equation 6.2), allowing the nature of behaviour implied by a state to change gradually across session boundaries (Roy et al., 2021) (we call this the ‘slow process’). (iii) Whereas for HMMs, the numbers of trials for which a single state remains active always follows an exponential distribution, we adopt a semi-Markovian approach, allowing for more general distributions (which duly provided a better fit to the data). The prior over these duration distributions encourages temporally extended states, in order to extract persistent behavioural modes, rather than single trial deviations which are more likely noise. Taking all these additions together we end up with a dynamic infinite input-output hidden semi-Markov model (abbreviated as diHMM).

The transition matrix over a flexible number of states and the evolution of the psychometric weights are defined by priors, and the Bernoulli observation model provides a likelihood for each trial, allowing for approximate Bayesian inference (further details in Methods). We performed this via a Markov chain Monte-Carlo algorithm, namely Gibbs sampling. For a single animal, the entire response and feature data across all training sessions were fitted together (individuals were fitted entirely separately). Integrating across a number of Gibbs samples from multiple Markov-chains led to a set of behavioural states defined by their session-varying weights ***w***_*x,n*_ and duration distributions, as well as a hard assignment of every training trial onto one of these states (we discuss a way to estimate how strongly a trial is connected to its state in the Methods). While all other relevant random variables are specified hierarchically or ruled by vague priors, the variance allowing slow changes within states is set, as inference over this variable proved problematic. We revisit this parameter in the discussion.

## 3 Single animal fit

We visually summarise the model fit to an individual at the resolution of entire sessions as shown in **Fig. 2**. This particular animal exemplified many of the interesting properties that can be found across the wider population of trained mice. The inferred model contains eight states, but these states were generally active for only a small number of sessions, before being replaced by others. Thus, in a typical session, the mouse used only a small number of states (usually, the majority of trials in a session is explained by a single state). Later states generally represented more adept behaviour, though not exclusively. The mouse started out with state 1 that exhibited a flat psychometric function (PMF; far right of the plot), indicating that the animal did not take into account the relevant feature, the side of the sensory input that was presented on the screen. This particular state only explained the very first session, before being replaced by state 2, which also has a flat PMF, only notably shifted. This shift in bias was strong enough to warrant a new state (rather than using the slow process to change the existing state), but there is no evidence that the animal advanced in its understanding of the underlying task.

**Fig. 2.**
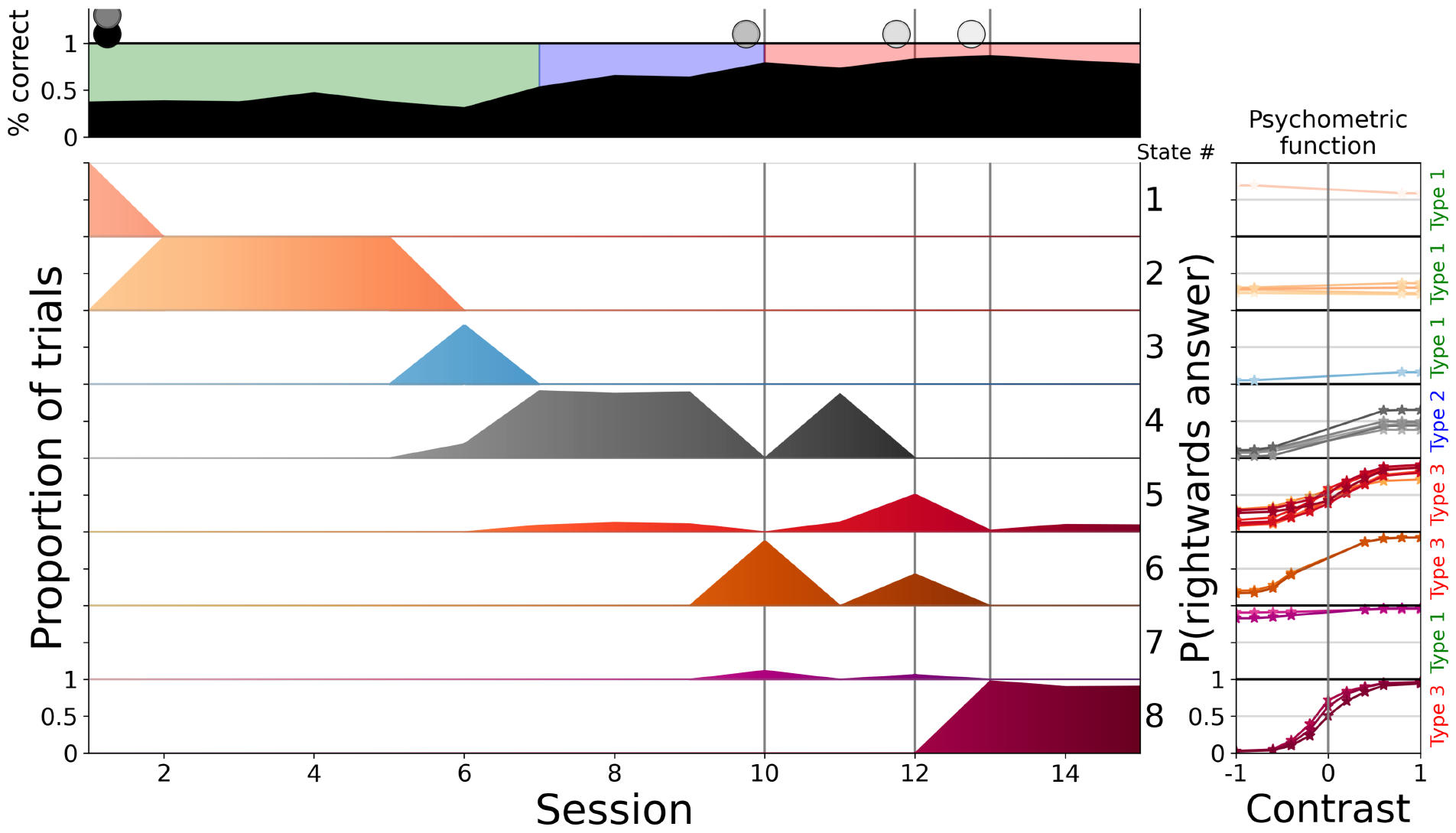
Dynamic infinite hidden Markov model (diHMM) fit to mouse KS014. The topmost row shows the overall performance during the session, as percent correct, and the current stage of learning as the background colour (we elaborate on learning stages later in the text). Vertical lines with shaded circles at the top indicate the sessions during which new contrasts were introduced. The remaining rows show the prevalent behavioural states (label to the right) ordered by appearance, indicating which percentage of trials they explained during each session. To the far right of every state we show its psychometric functions (PMFs) across time, ignoring the contribution from the history of previous choices. The saturation of the colours of the states indicates successive appearance, and match the PMF plots.

State 2 lasted for four sessions, meaning behaviour stayed relatively consistent during this time, before being predominantly replaced by state 3, with yet another flat PMF, which showed extremely biased behaviour (leading to an even lower reward rate, due to the bias correction). However, in the same session as the appearance of this state, we also finally saw the introduction of state 4 with a non-flat PMF. This new state showed very good performance on one side (the leftwards side in this case), and more or less random performance on the other side. It seems that the mouse was only taking sensory information from one side of the screen into account when making its choice, and performed randomly if that side was uninformative. It is important to keep in mind that such random behaviour for the other side was more rewarding than always giving the same answer. This is partly since this random answer will sometimes have been correct, by chance, and partly because the bias correction foiled perseverative behaviour by repeating mistaken trials. Thus, random behaviour would lead to more trials on the side at which the animal was proficient (though it would indeed have been better to choose the opposite side deterministically if the stimulus did not appear on the attended side). Along with state 4 we also saw the introduction of state 5, describing the behaviour at the ends of sessions 7, 8, 9, and 11 (and later also the ends of session 14 and 15). Puzzlingly, this state had a good PMF on both sides and a higher reward rate than state 4, but even though this better state was available, the animal seemed incapable or unwilling to use it for the majority of a session.

The “one-sided” state 4 remained active for some sessions, over the course of which the chance-performance side of the PMF gradually improved through the slow process, as can be seen in the evolving PMF (with darker colours showing later sessions). The last major step in learning however appeared abruptly again, as the animal showed a state (6) in which its performance was good on both sides (notably, this new behaviour was yet again different from state 5, though they both had the quality of being good on both sides). Together with state 6 we also saw the introduction of state 7, which captured a strong but transient decline in the quality of behaviour during sessions. Lastly, state 8 represented another notable change in behaviour, as the performance on 100% contrasts approached perfection sufficiently abruptly to warrant a new state, which allowed the mouse to conclude this part of training soon thereafter. Note that this model lacks a formal lapse or trembling ‘hand’ process, but it captures errors on easy contrasts by setting the psychometric weights of a state such that the PMF asymptotes at that error rate for the given range of contrasts.

Our model also affords a fine-grained look at the use of behavioural states within a session. Although the diHMM provides a full posterior over the states for each trial, this is not directly useful, due to technicalities of the sampling procedure. We therefore processed the chains of samples to extract a measure of how much a trial belonged to a state (details in Methods). We show an excerpt of this for session 12 of the animal we discussed so far in **Fig. 3**. This shows two rather clear transitions between states. The reasons for the animal to have made such a transition are probably multi-faceted, and may have been both internal, such as insights into the task leading to changed behaviour and more rewards, high or low motivation to obtain reward (Berditchevskaia, Cazé, & Schultz, 2016), as well as external, such as for example a number of low contrast, perchance unrewarded trials serving to demotivate the animal. We do not model these reasons, and instead only describe observed changes.

**Fig. 3.**
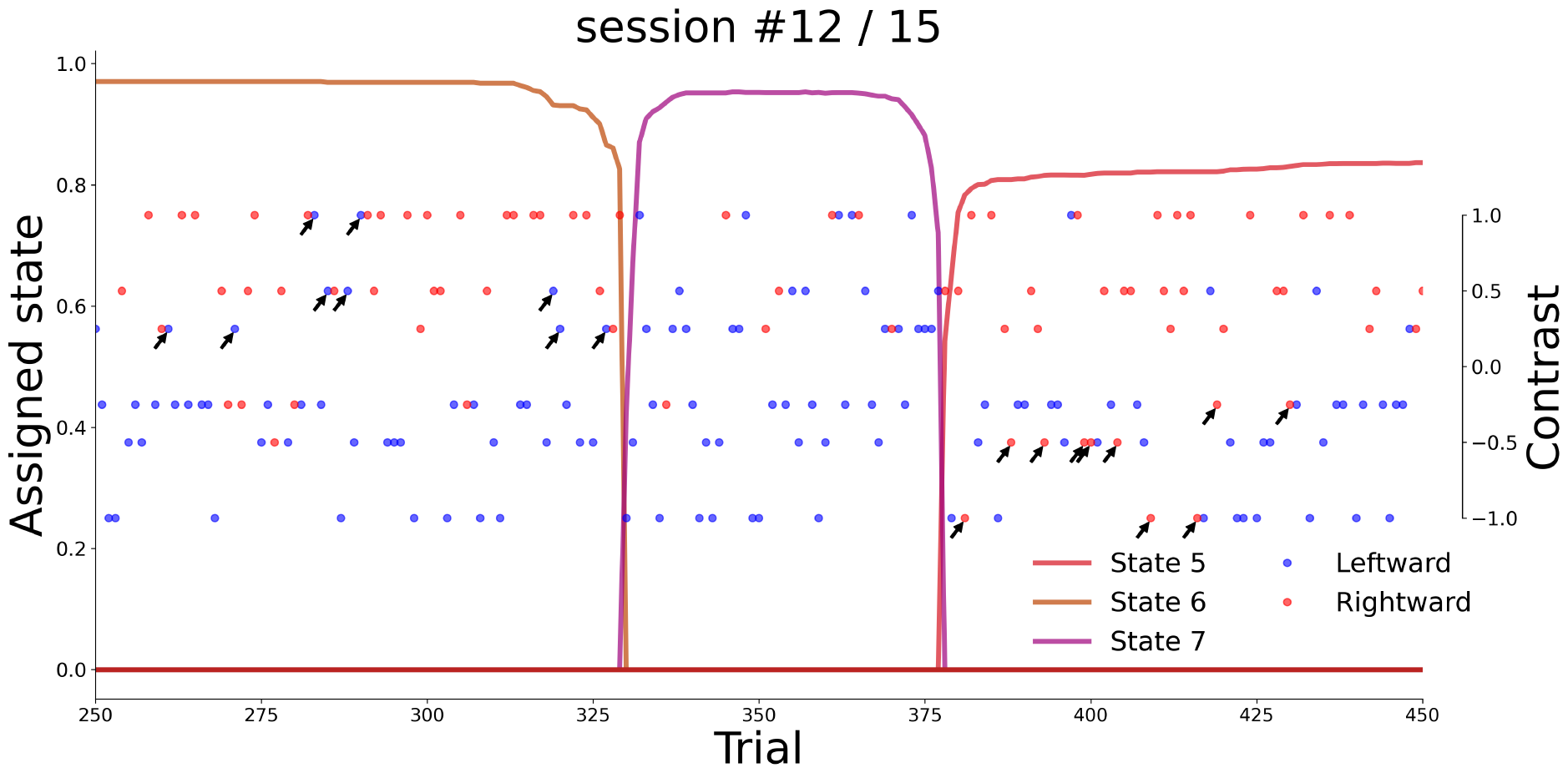
Excerpt of state assignments in session #12 from the mouse of **Fig. 2**. The left y-axis serves as a scale for how connected a trial is to the other trials of that state (see Methods for details). The right y-axis shows the contrast. The dot colour indicates the animal’s response. One can see how the drastic and sudden change in the response patterns, leftwards (blue) answers for rightwards (positive) contrasts, from trial *∼*330 to *∼*380 is detected by the model with a state transition. The PMFs of state 5 and 6 looked similar, but did in fact represent significantly higher rates of errors on the right and left side respectively, we highlighted these mistakes with arrows.

The within-session fit showcases two interesting points about the model. First, it is able to find temporary, but strong deviations in behaviour. State 7 only explained a couple of dozen trials in two sessions, but while the animal was in this state, it acted in an extremely biased way (comparable, but flipped relative to its earlier state 4, with the notable difference that state 7 was much briefer in duration when it appeared, and we know that this state could not have originated from a lack of task understanding, but was presumably some form of inattention). This change in behaviour can be spotted directly in the response patterns of the animal, and different Gibbs samples of the model’s posterior consistently used a separate state to characterize the trials concerned.

The second point is more subtle, and less directly observable in the mouse’s responses: the model used different states to explain behaviour before and after the biased trials explained by state 7, even though behaviour looked about equally good in both time periods. Going into the details of the fit, we can see that the model assigned different error rates on easy contrasts to the two states, and this can be found in the choices: the proportions of choices for the two different sides are significantly different for the two states (ANOVA for responses during trials of state 5 or 6 in session 12, with factors signed contrast and state, the latter having two levels, state 5 and state 6: the factor state is significant with p=0.0011), which can also be gleaned from response dots marked by arrows in those periods in **Fig. 3)**.

## 4 Fits across the population

The three-fold progression we observed throughout learning in **Fig. 2**, from flat PMFs, to “one-sided” behaviour, to generally good performance, is typical for the population of mice we fitted. To define this more objectively, we clustered the states into these three types, based on their reward rate, i.e. for every state we took its PMF (ignoring perseveration) and computed the expected number of correct responses on easy trials (100% or 50% contrast, since some PMFs were only well defined for these contrasts, additionally, including more difficult contrasts can lead to lower reward rates for more broadly defined PMFs even though they are better on easy contrasts). The boundary between type 1 and 2 is at 60% reward rate, and between type 2 and 3 at 78% reward rate (details in Methods). We show an overview and some examples of the different types in **Fig. 4**. Note that this only exhibits the first PMF of each state, i.e. the behaviour it described at its inception (as states could change their type via the slow process), thereby representing response characteristics after a notable discontinuity in behaviour.

**Fig. 4.**
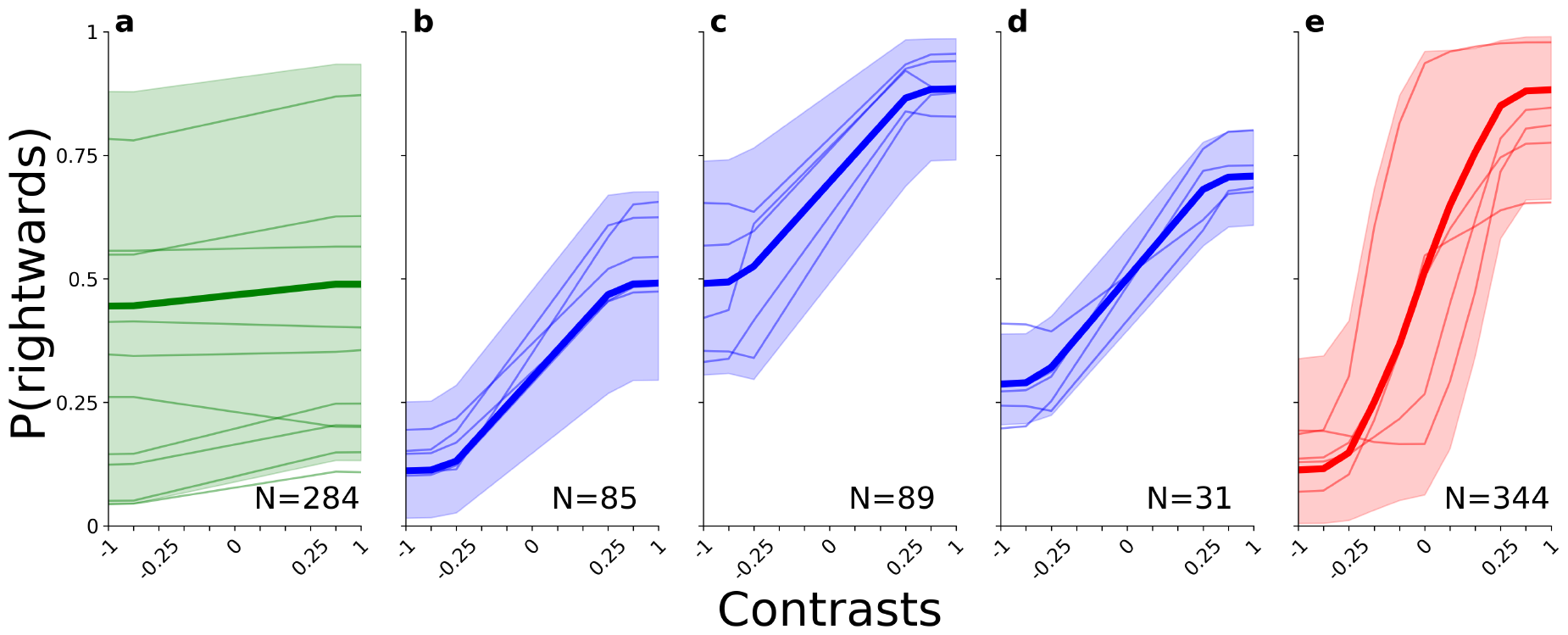
Summary of the PMFs associated with the different types, for this we use the first PMF of every state in every animal. Each subplot shows a specific type: (a) type 1 in green; (b-d) type 2 in blue, further split by whether the PMF is left-biased (b), right-biased (c), or symmetric (d); and (e) type 3 in red. The thick lines indicate the overall mean over PMFs of the type, the shaded regions show the range in which 95% of the PMFs fell (computed separately for each contrast level), and the thin lines show samples of individual PMFs of these groups. Text in the bottom right indicates how many PMFs of this type were present across the entire population.

In addition to the state types, we define the *stage* at which an animal is, as the highest type it has so far used for the majority of any previous session. For instance, if up to session *s−*1, an animal only used type 1 states, or type 2 states for fewer than 50% of trials, then it would be in stage 1 for those sessions. If on session *s*, it then used type 2 states for more than 50% of trials, it would switch to stage 2 on that session. Since the state types delineate structurally different aspects of task understanding, the stages allow us to determine for how many sessions the animals stayed at a certain level of understanding. Stage 1 involves a sequence of one or more states with flat PMFs which could show a whole range of biases, but share the property that the contrast location seemed not to be taken into account to determine choices. Stage 2 almost always involves states that showed good performance for stimuli on either the right or the left side, but close to uniform guessing for the other side. Only rarely were intermediate PMFs nearly equally good on both sides, **Fig. 4**b and c account for 85% of intermediate PMFs, and d for the remaining 15% (those rare cases were still closer to other type 2 states than type 3, which we verified by looking at their appearance time during training). After some time in such partially sensitive states, the animals started, in stage 3, apparently paying attention to both sides. The more difficult contrasts were usually quickly introduced once such states were reached, as they tended to be relatively symmetric, and therefore the type requirement for this stage (a close to 80% reward rate or higher), also led to a fulfillment of the criterion for contrast introductions. Generally, it took some further refinement of initial type 3 states, through the reduction of errors on easy trials on either side, to master this stage of training and progress to the next level of shaping in the IBL task.

The progression through state types was not monotonic, since multiple states of potentially different types could be simultaneously present in a session, and, e.g. a state of type 2 could dominate in a session after other states of type 3, as in **Fig. 2** on session 11. Nevertheless, the stage classification is monotonic, as it reports the higher level of performance which the animal is known to have reached.

The three stages segment the learning process. Firstly, we can analyse the proportion of training time the animals spent in the different stages, by showing these proportions on a simplex (**Fig. 5**). The classification into the types proves its behavioural relevance, as the large majority of animals needed some time in each of the stages (i.e. there are only few mice on the edges of the simplex). As we can see, most animals spent the longest time in stage 3 – that is to go from moderately competent performance (based on the prevalence of states that implied at least a reasonable understanding of the task), to behaviour that sufficed to pass the rather stringent training criteria. This was unexpected, as no fundamental change in understanding seems necessary, unlike the changes from stage 1 to 2 (where the animal has presumably to learn to pay attention to the Gabor patch in the first place); and from stage 2 to 3 (where it must learn to pay attention to both sides of the screen). However, reaching the required accuracy seemed difficult, even once the principles of the task were understood. Of course, the increase in reward rate was not great in this period, so there might have been less pressure to improve. Some of the very longest trajectories (the largest circles) were associated with especially large numbers of sessions in stage 3, but overall the average fractional occupation was remarkably consistent across training lengths (for a median split on the total training time, the mean relative occupancy for stage 1, 2, and 3 respectively are: shorter half (0.24, 0.17, 0.59), longer half (0.21, 0.15, 0.64)). Type 2 is the stage which consistently lasted for the fewest sessions, implying that the mice did manage to pay attention to both sides not too long after starting to pay attention to one side.

**Fig. 5.**
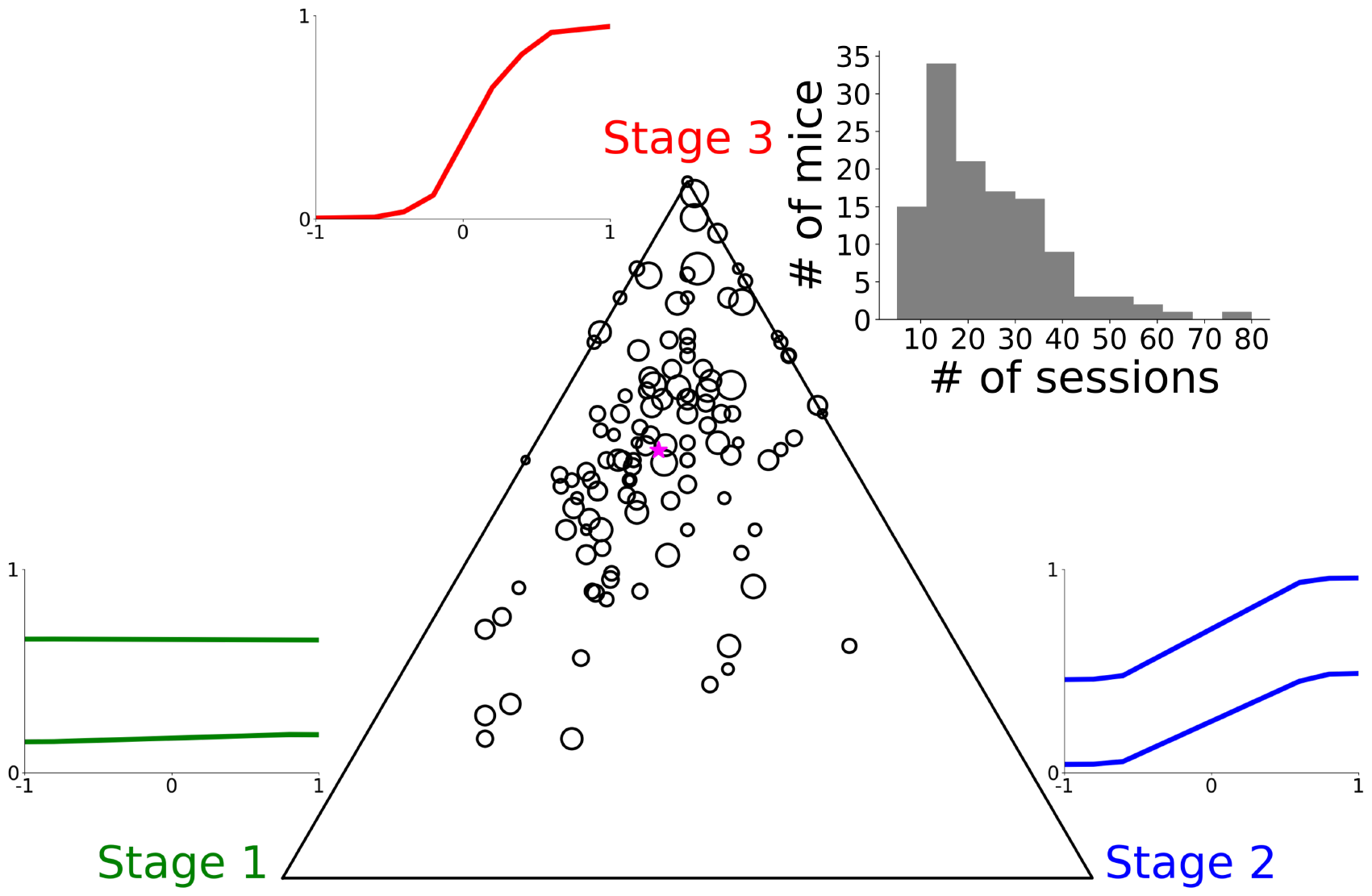
Proportions of sessions it took each mouse to reach the next major step in training, as defined by the 3 stages, scattered as circles on a simplex (the larger the proportion of sessions within a specific stage, the closer the dot for that animal is to that corner of the simplex). Simplex corners are identified by example PMFs from the type of that stage. Marker area indicates the total number of sessions (min=5, max=80). The magenta star marks the average proportion, the size of the star indicates the mean number of sessions (which was 24.2). The histogram shows the overall distribution over the number of training sessions.

Connected to this is the question of how the two kinds of change affect behaviour. For the slow process (gradual changes within a state), we analysed this by comparing the weights of the PMFs at the first session of a state, to those for its last session. For the fast process (new state introductions), we compared the weights of a newly introduced state to the closest previous state, as determined by the Wasserstein metric on their resulting PMFs (ignoring the perseverative weight). To highlight the change most clearly, we focussed on states which bring the animal into a new stage. These weight evolutions, split by the different types, can be seen in **Fig. 6**. As the main driver of performance, contrast sensitivities reliably increased both over the lifetime of a state and when a new state got introduced. Surprisingly however, we observed that both the bias and perseverative weights were remarkably stable within a state, even though these weights were not beneficial to performance. The bias tended to change more substantially upon the introduction of a new state, marking a decided difference between what the two types of change accomplished. When comparing the distributions of weight changes of the bias weights (see **Fig. A16** and **Fig. A17**), the changes through the fast process tended to be significantly larger (Mann-Whitney-U-test on absolute weight changes, 2 biases, 2 fast change points, 3 slow change processes, for 9 out of these 12 comparisons, the fast changes had a significantly larger change at a 0.05 significance level). We can also see that the perseveration weight played a small, but consistent role throughout learning (though its relative influence waned as the sensitivities kept growing).

**Fig. 6.**
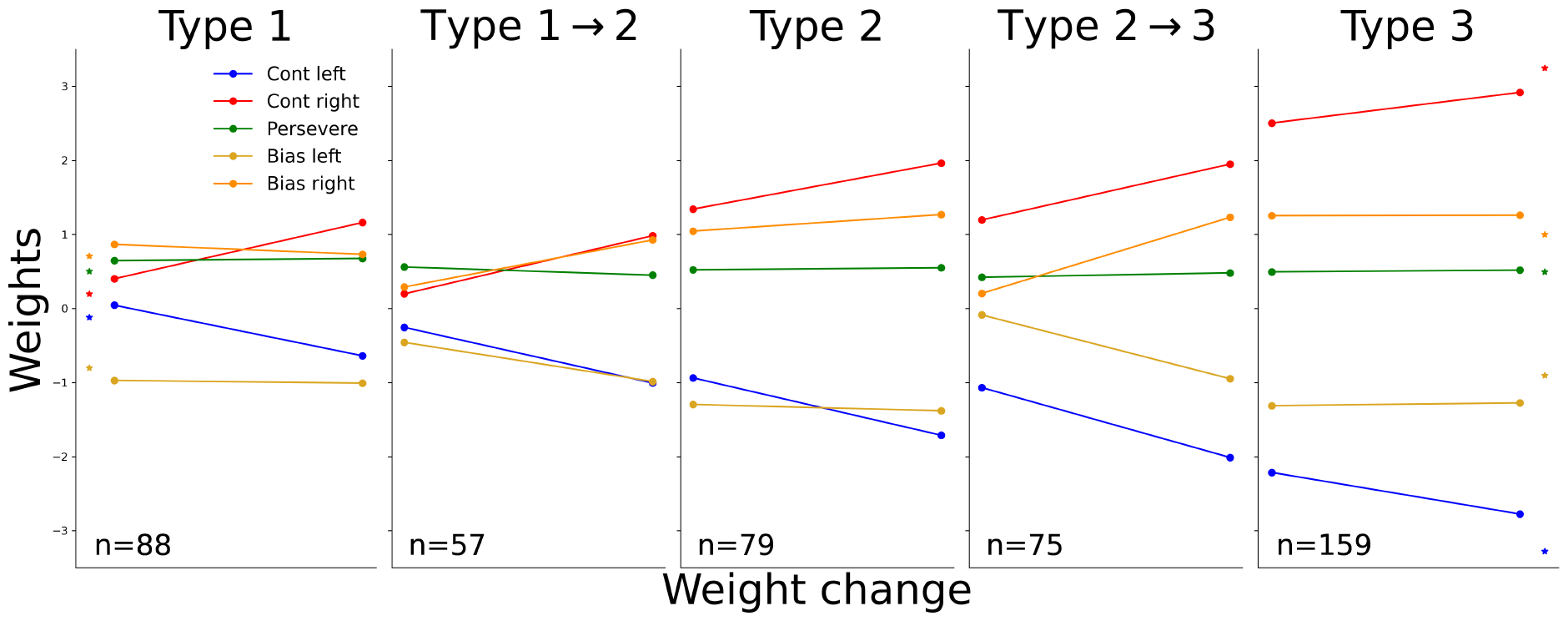
Evolution of the weights of states on average, through slow and sudden changes. Subplots titled by a type represent the weight changes from the first appearance of a state of this type to its last, so only state-internal slow changes (we only considered states which appear on at least 5 sessions, but not more than 15, as both extremes would skew these averages, since states might have changed their type in this time). Subplots with a title indicating a transition from one type to another show how much each weight of the new state differed from the weights of the closest previously existing state, and is based exclusively on the states which first brought the mouse into a new stage (i.e. for “Type 1→2”, we only took into account the first type 2 state exhibited by the mouse, and only when that state is type 2 from its inception). Coloured stars on the leftmost and rightmost plot indicate the average value of the weights of only the very first states of each mouse and the weights of the dominant state on the last session, respectively. We split the bias weights into left-biased and right-biased, as they would otherwise cancel out. Whereas contrast sensitivities increased both through fast and slow changes, it is noticeable that biases stayed almost constant throughout the lifetime of a state on average, but changed more noticeably through sudden transitions.

The introduction of new states signifies notable changes in behaviour, so by studying the patterns of their occurrences, we gain insight into when behaviour was volatile or when substantial progress was made. The two histograms in **Fig. 7** show when new states first appeared across normalised training and session time, for the entire population of mice. Of course, all animals needed to introduce at least one new state in the first session, to explain any behaviour, therefore we excluded this first state so as not to skew the histograms. In subsequent sessions, gradually fewer states tended to be introduced, indicating that behaviour saw fewer drastic changes as training progressed. We observed earlier that animals spent most of their time in stage 3, i.e. perfecting their behaviour, and we can now conclude that this mostly came about through gradual improvements of behaviour, rather than sudden marked changes. The pattern of introductions within individual sessions is even more striking: the majority of states were introduced at the very start of a session. This resonates with previous findings about changepoints in behaviour occurring at session boundaries (Papachristos & Gallistel, 2006). Apart from this strong trend, there seems to be a slight tendency for new states to get introduced towards the end of sessions, which might have partly been inattentive or demotivated states which the animals often fell into at the end of a session, and which the model sometimes picked up on when they were consistent and long enough (the end of a session is triggered if it is longer than 90 minutes, or due to slow response times).

**Fig. 7.**
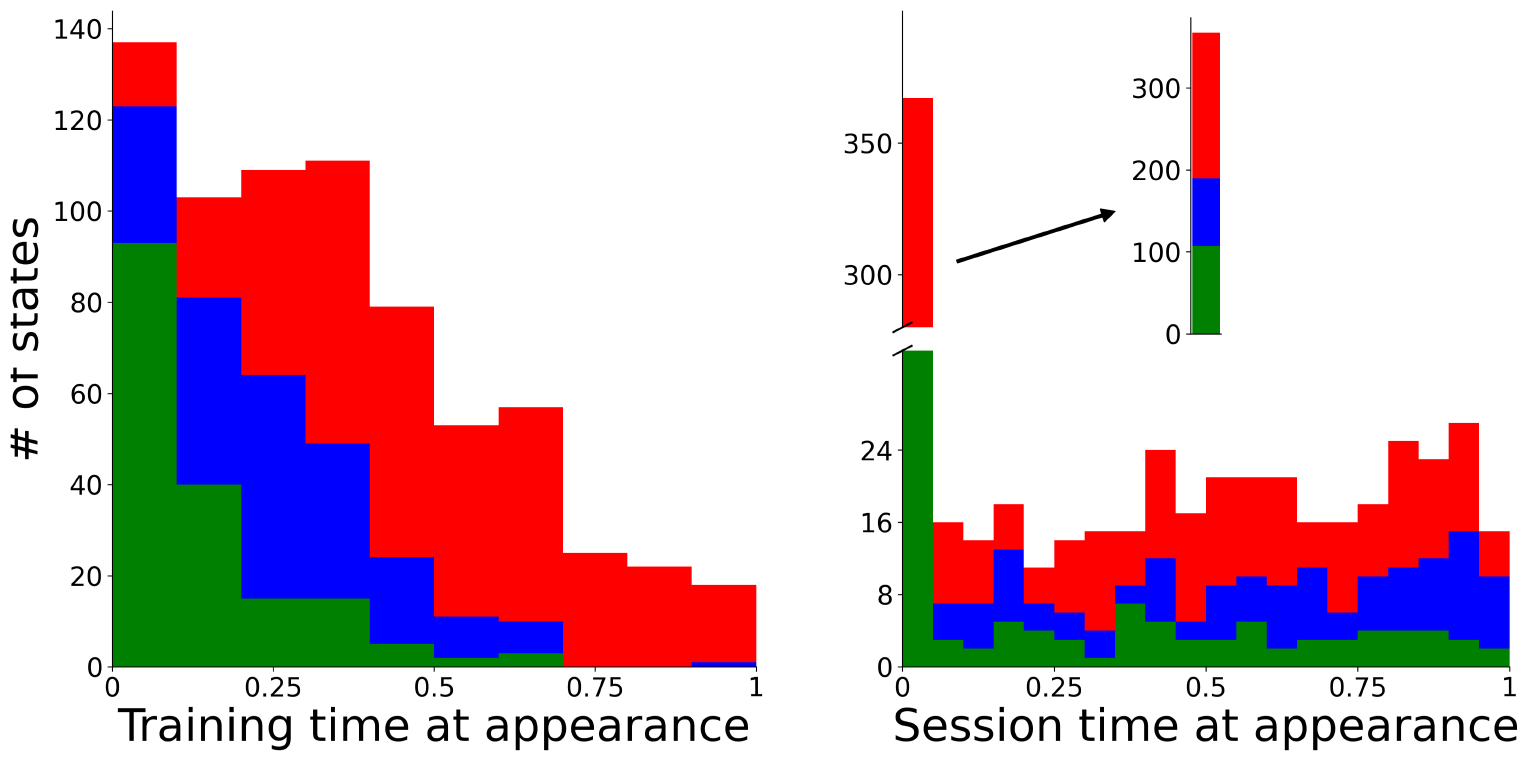
Histograms of all state introductions, excluding the first state of every animal, across all of training (left) and within sessions (right). We colour by state type and normalise the entire length of training of an animal, and all individual sessions, onto the range between 0 and 1 for comparison purposes. The inset on the right plot shows the bar of the first time bin uninterrupted.

### 4.1 Inter-individual differences and variability

So far, we have highlighted general patterns during learning, but perhaps even more salient than these similarities was the wide-ranging variability across animals. Inter-individual differences, especially during learning, are a known phenomenon (The International Brain Laboratory et al., 2021), though rather less commonly studied (though, for instance, see Akiti et al. (2022); Kastner et al. (2022)). Such differences are already visible in many of the plots above. Biases in type 1 states spanned the entire range of possible response patterns. Similarly, type 2 PMFs appear to have been randomly biased to one side or another, or, in rare cases, were symmetric and largely unbiased, but did not yet show good performance. We were particularly surprised to notice that we could find no regularity between type 1 and type 2 biases. Of the 57 mice in which type 2 onset occured suddenly, see **Fig. 6**, 32 had expressed the same direction of bias as the new type 2 state in any previous type 1 state, whereas 25 had not (two-sided Binomial test for whether the proportion of previously expressed biases differs from 0.5 gives p=0.427). Here, we define leftwards, rightwards, and no bias as whether the average rightwards response probability of a state’s PMF is below 0.45, above 0.55, or between those two values respectively. This means that one cannot predict future biases of the animal from its stage one biases.

The number of sessions mice required to learn also varied widely, spanning an entire order of magnitude. Similarly, the number of sessions spent in the different stages was also highly variable. To gain insight into the factors underlying the learning steps between the stages, we analysed the correlations between the number of sessions spent in them. The simplex plot does not strongly indicate any patterns, we quantify this by analysing their correlations to one another: duration of stage 1 to stage 2: Pearson’s r=0.31, p=0.0007; stage 1 to stage 3: Pearson’s r=-0.004, p=0.96; stage 2 to stage 3: Pearson’s r=0.27, p=0.0029. Notably, the main chunks of training time, stage 1 and 3, show no correlation whatsoever. A speedy understanding of the basic contingency of the task therefore did not necessarily go along with the ability (or will) to perfect this behaviour quickly, suggesting that they required different competences. The strongest correlation exists between stage 1 and 2, which makes sense in so far as they were both concerned with discovering how to make use of the stimulus information.

Another phenomenon complicating broad conclusions about the learning trajectory is the prevalence of sessions in which the animal did not use its current best state (in terms of reward rate) for the majority of trials. The existence of such sessions prevents us from viewing learning as a process of discovering new kinds of behaviour, and then using them based on their performance (as indicated by the amount of reward collected), which would make behaviour monotonically improving. To analyse these regressions, we considered the expected reward rate of the PMF of a state, keeping track of which states have been used so far and updating their performance as they changed slowly across time. We define a session to have regressed if the state which explained the majority of trials was more than 2.5% worse (corresponding to 5% of the total range, since reward rates were roughly limited to the range of 0.5 to 1) in terms of reward rate than the best state of previous sessions (for this, as for the state type classification, we computed an idealised reward rate between 0.5 and 1, not taking into account debiasing and non-strong contrasts). Such regressions occured multiple times in the example animal shown in **Fig. 2**: state 5, which was considerably better than state 4, was introduced towards the end of session 7; nevertheless state 4 was responsible for the majority of sessions 8 and 9, with state 5 again confined to the end of the sessions. Similarly, state 4 was used in session 11, even though both states 5 and 6 were better; and session 12 was dominated by state 5 which by then was worse than state 6.

Looking at such regressions across the population, we can see that they occured in more than 85% of the fitted animals and had a notable effect on the overall training time, see **Fig. 8**. We were unable to determine any causes for these regressions, in particular the time between the previous session and the current one had no significant correlation to the amount of reward lost through the regression (Pearson’s r=-0.006, p=0.78), and the set of times between the previous and a regressed session was not significantly different from the set of times between a previous and a non-regressed session (Mann-Whitney-U-test p=0.74).

**Fig. 8.**
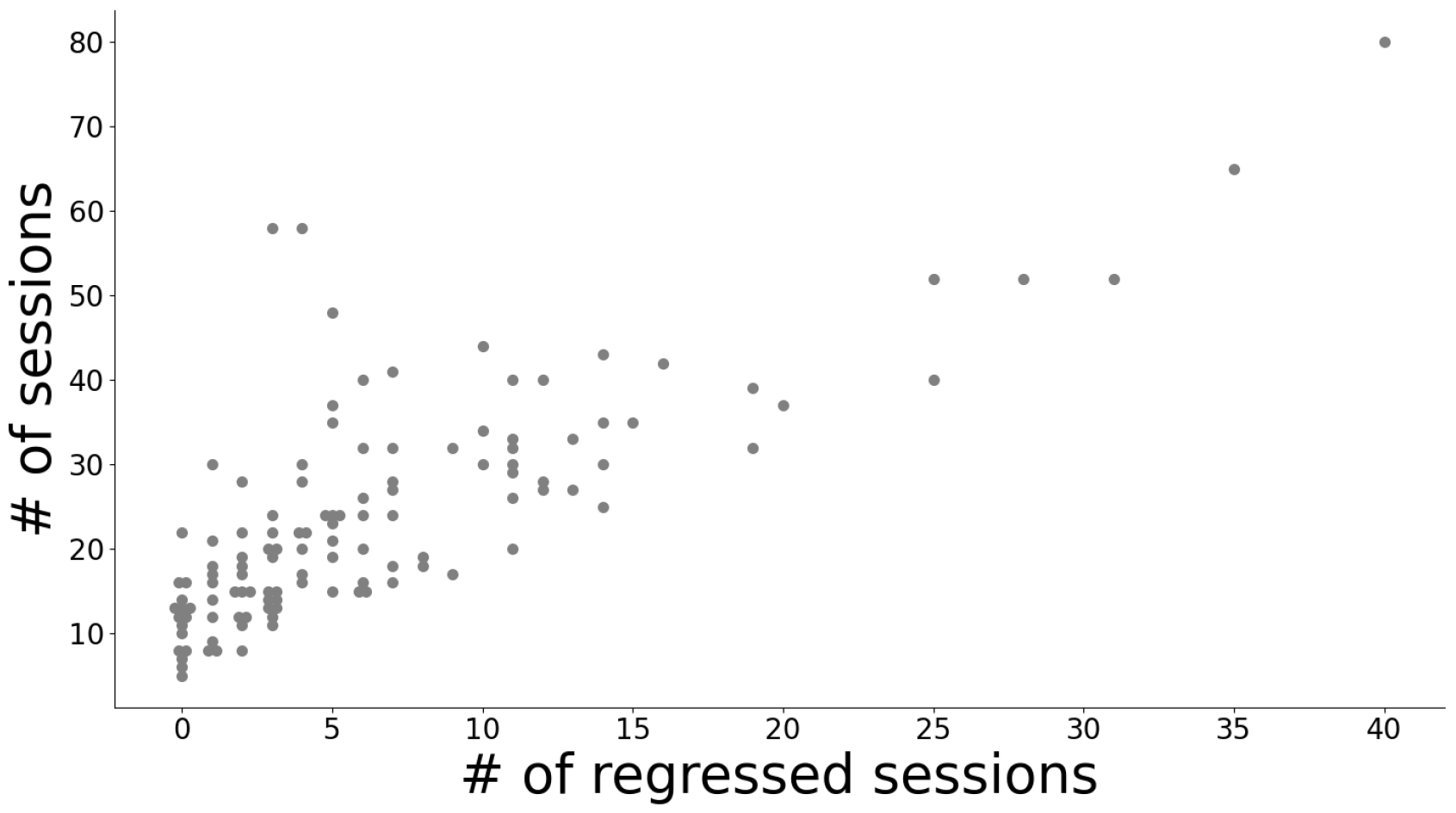
Scatterplot of the total number of sessions of an animal against the number of sessions in which a state with a worse PMF than the currently best available one explained the majority of trials. We add an offset to points coinciding at the same spot for visibility. While there is considerable variability for any given number of regressed sessions, there is a clear positive correlation (Pearson’s r=0.78, p*<* 1 *×* 10^*−*24^).

**Fig. 9.**
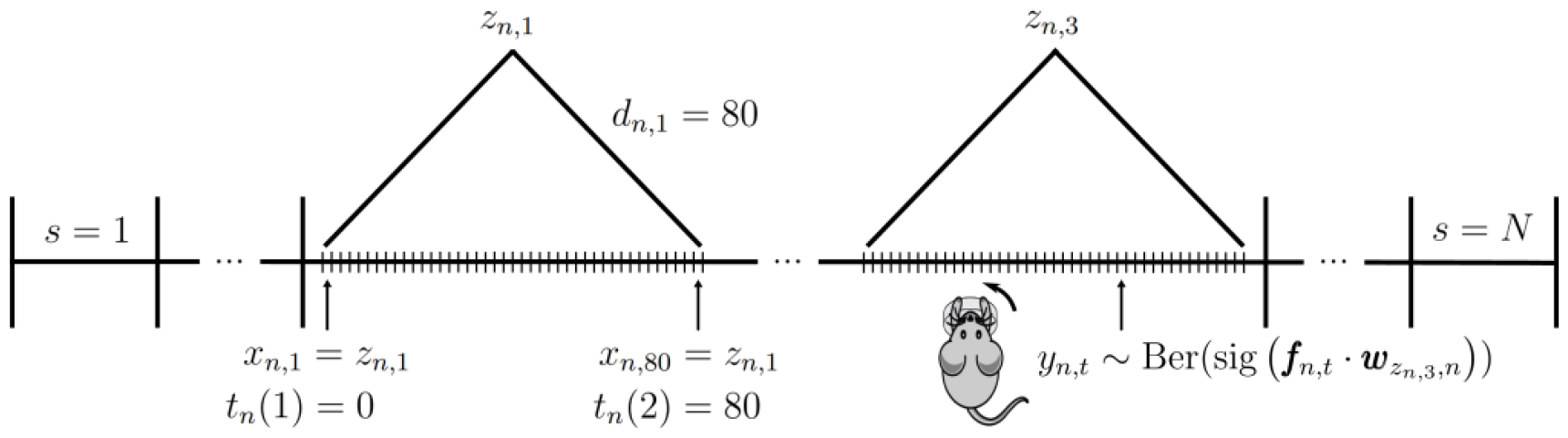
Visualisation of the different variables across training.

15% of mice never had a bad session in the sense of a worse reward rate, and these were generally quite fast learners. However, we also see that slow learners did not necessarily have many regressed sessions, as two mice with close to 60 sessions only had 4 and 5 regressed sessions, meaning their performance was mostly monotonically increasing.

## 5 Discussion

We have presented a highly flexible model which is designed to describe the stages of learning from the very first day an animal interacts with a task until it becomes an expert. Using it on the shaping sessions of the IBL decision-making task, we distinguished fast, abrupt transitions in behaviour, and slower, gradual ones. Learning on this task decomposed into three distinct stages, through which almost all animals went: Initial, undifferentiated, and often biased behaviour, partial, one-sided understanding of task contingencies, and lastly full understanding of the task. While these broad stroke characteristics were consistent across mice, the details of behaviour in these stages differed largely across the population. Similarly, the way they progressed through these stages differed widely in duration and composition of sudden and gradual steps. In addition to improvements through learning however, animals also often regressed in their performance for large parts of sessions, adding another layer of intricacy to the study of the learning trajectory.

We found only a small correlation between the time it took individual mice to progress through some of the behavioural stages, suggesting that they had to draw upon largely different skills to learn the requirements of the task. Similarly, animals expressed varying biases across the stages of learning, without notable tendencies to repeat previous biases. This leads us to speculate that there are different sources of biases: initially, in stage 1, when the mouse paid no attention to the stimulus, it might have been a motor bias, in stage 2 the bias could have been an expression of which side the animal happened to notice first as being informative, and in stage 3 the bias might have stemmed from differences in sensory acuity. In the IBL training scheme, after the sessions we analysed, the mice underwent a further phase (‘biased block training’, in which left or right stimuli dominated in blocks of 20-100 trials). Consistent with our other results, the length of this phase also turns out not to correlate with the total pre-bias duration, nor to any of the stage durations (see section 6.5 for details). This shows yet again that learning was dominated by a large number of factors, and is seemingly difficult to predict.

Our modelling approach has a number of limitations. First, the setting of the slow change variance parameter which determined how much the behaviour of a state could change from one session to the next plays a critical role in steering the trade-off between introducing a new state versus adapting an existing one. We optimised this parameter in terms of cross-validation performance for the entire population, under the limitation that it be small enough that states were independently meaningful. However, the magnitude of slow changes may well depend on the individual, or might even vary across training time, and thus a more differentiated treatment might be appropriate. Furthermore, slow changes may also occur within a session (Roy et al., 2021), which could be incorporated into the model by adding additional time points at which weights can change. Another desirable extension would be to allow the duration distributions also to change over sessions. As training progresses, an animal might e.g. be able to use a high performant state for longer, which could be reflected via a changing duration distribution.

The model may also be extended by adding more observations for the states to explain, as the binary choice behaviour may limit the power to distinguish different behavioural modes. One obvious possibility are the reaction times of the animal’s choices; in principle, this would only require adding a suitable distribution to produce times for each state (e.g., from a drift diffusion decision-making process; Gold and Shadlen (2002); Ratcliff and McKoon (2008)). It would likely be necessary to make the distributions dynamic (by making some of the parameters of the drift-diffusion model dynamic), as the reaction times will surely improve with training, and may do so in quite rapid ways. Beyond this, one could add details of mouse posture to the states, such as pupil dilation and bespoke aspects of body posture, though this would require well-tailored distributions to describe (Wiltschko et al., 2015).

Besides learning, our approach to capturing behavioural evolution should be well suited to model other progressive changes, such as those occurring during ageing (Nyberg, Lövdén, Riklund, Lindenberger, & Bäckman, 2012). Fast and slow transitions between stages have also been studied for the acquisition of movement sequences (Luft & Buitrago, 2005). In this context, changes which occur between sessions are labelled slow, i.e. improvement through consolidation during breaks. Improvements during a session are considered fast changes. In our work we divide fast and slow changes via the magnitude of the change in psychometric weights, but only allow fast changes during a session (the slow process can only change weights at session breaks), and capture phenomena like an initial warm-up period through the usage of multiple states. Our results contrast with those in motor skill learning, in that if we examine the new states which bring the animals forward substantially, the vast majority occur for the first time at the start of a session, i.e., through what this literature would label as slow learning.

Of particular note are the rather long training trajectories. One might *a priori* assume that these are instantiated via the usage of a larger number of states, but this is not the case. Instead, we often see that few states are taken a long way via small steps, from uninformed to proficient (see **Fig. A15**). It will be important to assess the underlying nature of these states and their progression, by tracking neural data through the course of learning. Note that recovery analyses show that the model can cope effectively with such trajectories, without, for instance, inferring wantonly many states (see **Fig. 14**).

Apart from general insight into learning dynamics, our characterisation could help with the fine-tuning of shaping protocols to facilitate task acquisition. For instance. we observed that the animals initially need to learn to connect the stimulus side information with their behaviour, therefore, if quickly reaching expert behaviour is the only goal, it would seem ideal to make this association as salient as possible, for instance by making it dynamic (’wiggling’) in early trials. Conversely, having the more difficult contrasts present from the start might make the task much more difficult to learn, as the connection between actions and feedback would seem more probabilistic to the animals. We also saw that the last stage of training takes the longest time, in which the animal brings its performances to a consistent enough level to pass this stage of training. We found that the biggest contributor to this, i.e. the aspect of behaviour which held them back the longest (for the IBL-specific requirements), was insufficient consistency on easy contrasts. To accelerate this process it might be necessary to manipulate the reward rate, as the animals get a reasonably high reward rate without being too consistent. It might therefore be worth trying to only reward the animal after two correct responses in a row by the time it gets to stage 3.

Previous work using an HMM-based approach discovered demotivated states in behaviour (Ashwood et al., 2022), in a component of the IBL task that happened after the learning trials that we modelled. While we do occasionally find these (as discussed in the first example mouse), they were more rare than might have been expected from that work. For us, a majority of sessions were explained by just a single state. The dominant source of behavioural variability in our data came from learning and other large jumps in psychometric space, therefore the model used its capacity to capture these, rather than more subtle variations of proficient behaviour which can be found later. Especially towards the end of sessions, one can see a dip in performance of the animals purely by looking at the reward rate, on the order of tens of trials, which the model sometimes acknowledged with a separate state (seen in **Fig. 2** on a number of sessions). However, occasionally we just see a decrease in the prevalence of all states (meaning that these trials did not consistently enough get assigned the same state in the underlying Gibbs samples, see Methods). We interpret this as behaviour having been too variable during this period, and not reliable for long enough (even across multiple sessions) to have justified its own particular state in the model.

Our framework can be flexibly adapted to other cases of long-run learning. For instance, it is possible to tune the model to capture minute changes within sessions as opposed to broad stroke states across sessions, as here, by adjusting the propensity to infer new states for small changes in behaviour. Equally, the modular resampling procedure of the model allows it to be adapted to different kinds of observations, e.g. multinomial or Gaussian, by simply swapping out the inference mechanism of this one component of the model (although only some distributions are convenient for the gradual dynamics). We therefore hope that the tool we developed here will enable a wide range of researchers to study the development of behaviour in a systematic and revealing manner.

## 6 Methods

We structure these methods as follows: We begin with a detailed description of the infinite hidden semi-Markov model, i.e. the specification of the priors and how they relate to the other random variables in the hierarchy. In section 2 we describe inference for the logistic regression observation distributions, with a focus on the details of resampling, as these are not described in detail elsewhere. These two components together give us the full dynamic infinite input-output hidden semi-Markov model (diHMM). We cover how to make sense of the generated collection of samples in section 6.3, dealing with the hurdles of label switching and multi-modality, ultimately obtaining well-defined states from the samples. This concludes the details of the model itself. Section 6.4 elaborates how we assign states and their PMFs to the three types. Extending slightly beyond the scope of this work, section 6.5 relates properties of the initial learning considered here with the next training step, the biased block training. We conclude the methods by showing successful recoveries of various generative models in section 6.6.

### 6.1 Infinite hidden semi-Markov model

We start by describing the diHMM, focussing on Bayesian inference over its random variables. Following Johnson and Willsky (2013), we use Gibbs sampling, a Markov chain Monte-Carlo algorithm (MCMC), to realise an iterative resampling scheme over the model components, including the PMFs of the hidden states and the assignments of the individual trials onto those states. For this purpose, all distributions are paired up with conjugate priors in this section, to enable simple resampling steps. The posterior distribution is ultimately represented by a collection of samples, with every component being assigned an explicit value in each sample.

| Overview over quantities and counters |  |  |
| --- | --- | --- |
| Quantity | Meaning | Counter |
| N | number of sessions | $n$ |
| $T_n$ | number of trials within a session $n$ | $t, m$ |
| J | number of MCMC samples | $j$ |
| L | number of states in the model | $i$ |
| S | number of states employed within a session | $s$ |

We first describe all the relevant random variables (using the iterator notation from the table above).

The technical backbone of an infinite HMM is an hierarchical Dirichlet process. At the top of the hierarchy of this process is the prototypical transition vector

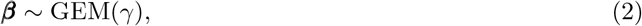

where GEM (named after Griffiths, Engen, and McCloskey) is a Dirichlet process without a base distribution, a pure stick-breaking process which samples a probability vector over infinitely many elements (which will be states in our case). The concentration parameter *γ* probabilistically determines the size of the individual sticks, and therefore how many states are practically relevant, with a higher *γ* allowing for more states. We put a vague Gamma prior on *γ*, making it, and thereby the propensity to introduce new states, part of the inference as well, with *γ ∼* Gamma(0.001, 0.001).

At the next level we sample the transition vectors, a classical HMM component, *π*_*i*_ of the individual states *i*. These are tied together via ***β***, which is used as the base distribution for a second Dirichlet process

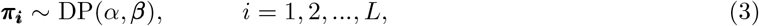

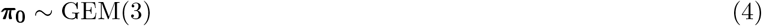

*α* is another concentration parameter and determines how closely the *π*_*i*_ are related to ***β***. Sampling the individual state transition vectors from this common source formalises an overall kind of state popularity. The higher *α*, the more like ***β*** is ***π***_***i***_, *∀i*, and so the more the bias in the frequency of state *i*^*′*^ in the particular sample ***β*** will be reflected in the transitions from *i* to *i*^*′*^, and so the more popular *i*^*′*^ will be overall. We put another vague Gamma prior on it, *α ∼* Gamma(0.1, 0.1). The initial state distribution ***π***_**0**_ is drawn entirely separately, with a concentration of 3 as a trade-off between allowing new states but not encouraging the invention of new states at the start of sessions.

For our inference scheme, we make use of the weak-limit approximation which puts an upper limit *L* = 15 on the number of states rather than employing the full infinite process. This simplifies the resampling scheme, while still behaving similarly to an infinite HMM if *L* is sufficiently large. Across the entire population, there is only one mouse with 15, one with 14, and three with 12 states; all other mice use fewer states. Furthermore, the minimum fraction of trials captured in states (as described further below) is 99.35% (mean is 99.996%), justifying the choice of *L* = 15. In particular, we still perform inference over the realized state complexity. In the weak-limit framework, Eq. 2, 3, and 4 turn into L-dimensional Dirichlet distributions

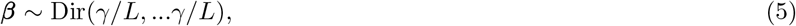

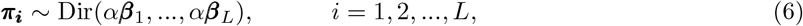

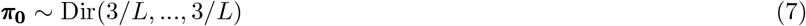

The transition structure within a session is given by

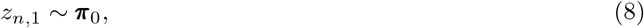

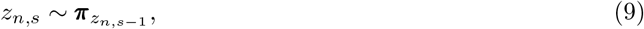

where *z*_*n,s*_ *∈ {*1 … *L}* is an indicator for the *s*th state within a session *n* (which does not align with the trial number), and *π*_0_ is the initial state distribution.

Given the transition vectors, the workings of the hidden semi-Markov model are fairly standard, except that the duration distributions are specified explicitly rather than being drawn from an exponential distribution (as in a regular HMM). We therefore also prohibit self-transitions, which makes a data augmentation scheme for resampling necessary, as described in Johnson and Willsky (2013). Nevertheless, as in a standard HMM, durations are statistically independent of the target state of transitions. Durations are drawn from a negative-binomial distribution, with state-specific random variables, coming from their own priors

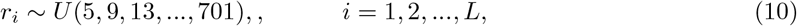

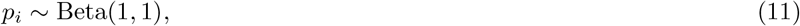

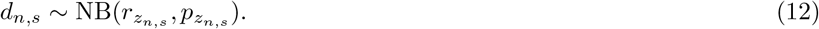

(Note the difference between state names *i*, which hold for the entire model, and the session specific state counters *s*, which can be used to find the current state name via the indicator *z*_*n,s*_). We chose a uniform prior over a large range of numbers for the possible values of *r*, to enable long durations, but excluded small values for *r* (in particular *r* = 1 would give the geometric distribution). Small values of *r* encourage transitions after a very small number of trials, which would capture the statistics of the presentation of left and right stimuli by the experimenter rather than the longer-lasting states that we sought.

States stay active and generate observations for as long as the drawn duration indicates

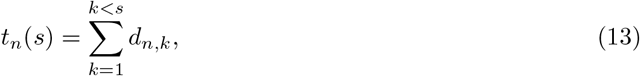

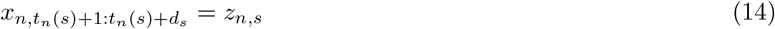

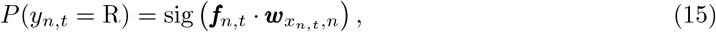

where we defined *t*_*n*_(*s*) to return the trial on which the *s*th state of a session *n* starts, which allows for the definition of *x*_*n,t*_, the state on any given trial *t*. We denote the logistic sigmoid function as sig. This takes the dot product between the state weights ***w***_*s,n*_ (which we discuss in the next section) and the input features of the current trial ***f***_*n,t*_ and produces the probability over the observation *y*_*n,t*_. The binary response variable *y* has 0 representing a leftwards, and 1 a rightwards, choice.

We summarise this collection of variables as

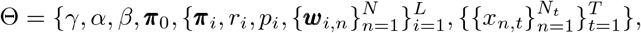

where *N* is the total number of sessions. The result of inference is a set of samples 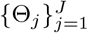. Each sample is a full instantiation of the listed random variables, which we can treat as a representation of the posterior. Gibbs sampling works by iterating through all variables, and re-sampling them from their distribution, given all other variables of the model. After updating all variables, the result is one new sample within the MCMC-chain. The details of how to resample the individual components can be found in (Johnson & Willsky, 2013).

### 6.2 Dynamic logistic regression prior and sampling

Gibbs sampling resamples each random variable conditioned on all others. Thus, inference over the observation distributions of the states is separate from almost all the rest of the model, only using the information of which trial is currently assigned to which state. We drop the explicit state dependence *i* in *w*_*i,t*_ for this section, but it is important to keep in mind that this sampling scheme is applied to every state individually, with each state *s* being influenced only by trials for which *x*_*n,t*_ = *s*. We implement slow changes in the characteristics of the states by putting a Gaussian random walk prior on the weights ***w***_*n*_, allowing for modest change across session boundaries, parameterised by the variance *σ*. We choose a diffuse initial distribution for the weights, and use cross-validation to select the inter-session variance *σ* = 0.03 (we performed cross-validation on a constrained range of small values, in order to limit the state adaptation process to small changes):

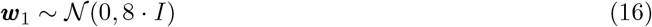

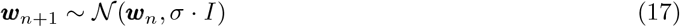

If a state has no trial assigned to it in a particular session, its weights are held fixed during the next transition, preventing states from morphing radically during a prolonged absence.

Inference for the logistic regression weights is performed using Pólya-Gamma data augmentation, which allows for efficient inference in settings with binomial likelihoods (Linderman, Johnson, & Adams, 2015; Polson, Scott, & Windle, 2013), since it is not possible to choose a conjugate prior. We review the relevant computations here, for a full treatment we refer to (Windle, Carvalho, Scott, & Sun, 2013). In the first step of the resampling scheme we sample pseudo-observations. This uses a Pólya-Gamma distribution PG, by first sampling *ω*_*n*_ *∼* PG(*b*_*n*_, ***ψ***_*n*_), where ***ψ***_*n*_ = ***f*** _*n*_ · ***w***_*n*_ is the dot product of features and weights, and *b*_*n*_ is the total number of times this exact instantiation of features was observed in session *n*. However, the same state is associated with more than just one specific instantiation of features (i.e., including contrasts of different strengths and side, and different response histories). To handle this, we treat a single session as multiple different time points, but prevent weight changes between time points that belong to the same session. In this way the observations from different features within the same session are effectively aggregated. To complete the pseudo-observation generation, we need *κ*_*n*_ = *a*_*n*_ *− b*_*n*_*/*2, where *a*_*n*_ is the number of rightwards answers observed for the current ***ψ***_*n*_ under consideration. Now *z*_*n*_ = *κ*_*n*_*/ω*_*n*_ can be treated as if they were drawn from *𝒩*(***ψ***_*n*_, 1*/ω*_*n*_).

This data-augmentation serves the purpose of having the ***w***_*n*_ emit observations with Gaussian noise (after combination with the features ***x***_*n*_ into ***ψ***_*n*_). Since the prior on ***w*** is a Gaussian random walk, this places inference in the well studied realm of Kalman filtering. To resample the ***w***_*n*_ we use the forward filter backward sample algorithm (FFBS, Carter and Kohn (1994); Frühwirth-Schnatter (1994)), which filters forwards through all the observations using a Kalman filter, then samples the sequence of ***w***_*n*_ backwards through time. A single resampling step therefore consists of first drawing the Pólya-Gamma variables to create pseudo-observations, then using them to sample the ***w***_*n*_ using the FFBS algorithm.

We consider four features for the logistic regression: the contrast on the left side, the contrast on the right side, an exponentially weighted history of five previous choices, and a bias. Separating the features for left and right contrast allows the sensitivities to the two sides to be different, an obvious property of mouse responses. Since the notional contrast values do not match the psychophysical difficulty of the contrasts (100% and 50% are both virtually equally easy to perceive, not a factor of 2 apart), we apply a transformation to have a better alignment. For this we follow Roy et al. (2021) and use a tanh-transformation, mapping the actual contrast *c* onto the input 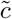 for our logistic regression through 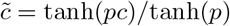, where we follow the recommendation and set *p* = 5, which scales the steepness of the transformation. This maps the contrasts (1, 0.5, 0.25, 0.125, 0.0625, 0) onto (1, 0.987, 0.848, 0.555, 0.302, 0).

The regressor for previous answers, enabling perseveration as a strategy, proved to be beneficial for cross-validated performance. It is associated with the famous law of exercise (Gershman, 2020; Thorndike, 1911), and has also been found to be exhibited by the mice in the asypmtotic regime that arises after the sessions that we are presently analysing (Findling et al., 2023). By contrast, there was no statistical support for a regressor sensitive to the interaction between past choice and past reward, as would be reflected, for instance, in win-stay, lose-shift behaviour. We implement the perseveration regressor as an exponentially weighted sum over the last few trials. We found that weighting the last 5 trials with an exponentially decaying filter with smoothing factor 0.3 worked best (though slightly different parameter settings have almost equal cross validation performance). Thus, we compute this feature on session *n* and trial *m* as such:

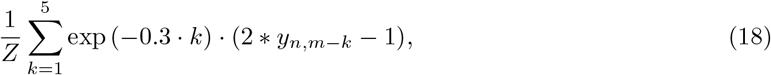

where 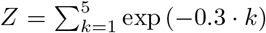 is a normalisation constant, such that the entire exponential filter adds to 1. The transformation 2 \**y*− 1 serves to encode responses as -1 and 1, for the purpose of having the perseverative feature sway the current response appropriately. Therefore, this feature reaches its maximal value of 1 if all 5 previous responses were rightwards and -1 if they were all leftward, putting it on the same scale as the other features. For the first five trials of a session, not all these previous trials are available, so missing ones are encoded as 0 (and we leave the normalisation constant *Z* unchanged). Time-out trials, where the animal did not respond before 60 s have passed, while skipped for the logistic regression of responses, are taken into account for the previous answer regressor, also encoded as 0.

### 6.3 Aggregation and interpretation of chains

We generally generated 60,000 samples from each of 16 chains (with different starting points), discarding the first 4000 as burn-in. We assessed convergence of the chains using the classical measure 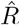 (Gelman & Rubin, 1992). This compares intra- and inter-chain variability of bespoke, state-independent features of the chains. To detect differences in the variances of the chains, and other problems which 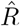 is known to miss, we also used folded-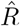 and rank-normalised-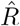 (Vehtari, Gelman, Simpson, Carpenter, & Bürkner, 2021). We reduced the memory cost by thinning the chain, using only every 50th sample (we did this purely for memory reasons, not because it is necessary for MCMC algorithms (Link & Eaton, 2012)). For a first pass, we sought to discard chains which differed substantially from other chains in the explored region in parameter space, either because they never reached the relevant parts of it, or because they spent disproportionate amounts of time in some modes over others. This is a known problem for MCMC algorithms in multi-modal environments, and can be mitigated by taking non-mixed chains and combining them via stacking (Yao, Vehtari, & Gelman, 2022). However, since our goal here is not prediction, we still want to focus on finding and visualising the most important modes of the posterior, which we did by combining the (possibly not perfectly mixed) chains, and considering the regions of probability space in which they collectively spent the most time. Given the slow transitions between different modes, we also did not split our individual MCMC chains when computing 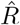, as the two halves of the chains were often too different.

As scalars underlying 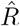, we used the concentration parameters: *α* and *γ*, as they are independent of states, and, as general properties of the fit: the number of trials assigned to the state with the most trials, and the second-most trials, as well as the overall numbers of states with more than 20%, and more than 10% of trials assigned to them (we chose multiple cutoffs to gain information about the fit at different levels of resolution). By greedily discarding the chains which increase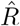 the most, we reduced the number of chains under consideration from 16 to at least 8. For this we considered all features and all variants of 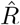(normal, folded, rank-normalised) at once, so we were minimising the maximum over all these 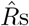. We only further processed the chains when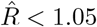, which is more conservative than some recommendations, but, in light of the strong multi-modality, more lenient than the newest ones (Vehtari et al., 2021).

However, it is still not trivial to extract information from the remaining chains given the multi-modality. There are two main sources of multi-modality: (i) genuine uncertainty in the usage of states or the exact setup of the random variables of the states, and (ii) mode equivalence with permuted labels (e.g., state *i* = 1 in the first chain might explain roughly the same set of trials as state *i* = 2 in the second). Although the second source makes evaluating the results more complicated, it is in fact just the sampling scheme working correctly, as there is nothing special about the particular state labels – solutions with permuted state labels are functionally equivalent. For the same reason, even within a single chain, a relatively consistent set of trials might be explained by one label for some part of the chain, but by a different label in another. Indeed, we frequently observed this kind of label switching, where one state completely took over the trials of another within a few sampling steps. In the limit of infinitely many samples, we can expect any trial to have a uniform distribution over the state label assigned to it; the only important question is which other trials were usually accounted for by the same state as the given trial within suitably similar samples.

To formalise the necessary abstraction from direct state assignments, we computed co-occupancy matrices *C*^*j*^ for each sample *j. C*^*j*^ is a matrix of size *T×T*, with *T* being the total number of trials across all sessions of a mouse, whose *t, m*^th^ entry reports whether trials *t* and *m* (for convenience, dropping the additional session label) used the same state in sample *j*

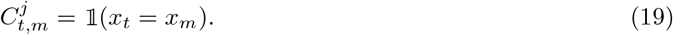

We used these co-occupancy matrices as a basis for two different processing steps: (i) at a coarser resolution across trials, we applied dimensionality reduction to find posterior modes; (ii) at full resolution, we averaged *C*^*j*^ across similar samples *j* to derive a matrix that describes the mutual affiliation of trials, allowing us to overcome the labelling issues. Both steps are reminiscent of representational similarity analysis (Kriegeskorte, Mur, & Bandettini, 2008), in that, instead of comparing two samples directly, we compare state co-occurrence within the samples.

In principle, to explore the posterior, we could have flattened each *C*^*j*^ into an *T* ^2^ vector and apply principal components analysis (PCA). However, there were too many trials per mouse (of the order of 13000) to do this at full resolution, so we binned the trials into 170 bins, ignoring session boundaries, and then used the Wasserstein distance to measure state co-occurence between the bins. That is, we define modified matrices *C*^*′j*^ as

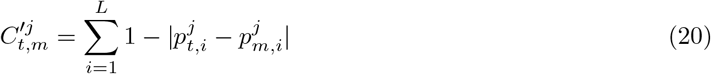

where 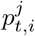 is the proportion of trials in bin *t* which is assigned to state *i* in sample *j. C*^*′j*^ reduces to *C*^*j*^ for bins comprising a single trial. We then plotted individual samples in the first three dimensions of the PCA-space arising from flattened versions of *C*^*′j*^, as shown in **Fig. 10**.

**Fig. 10.**
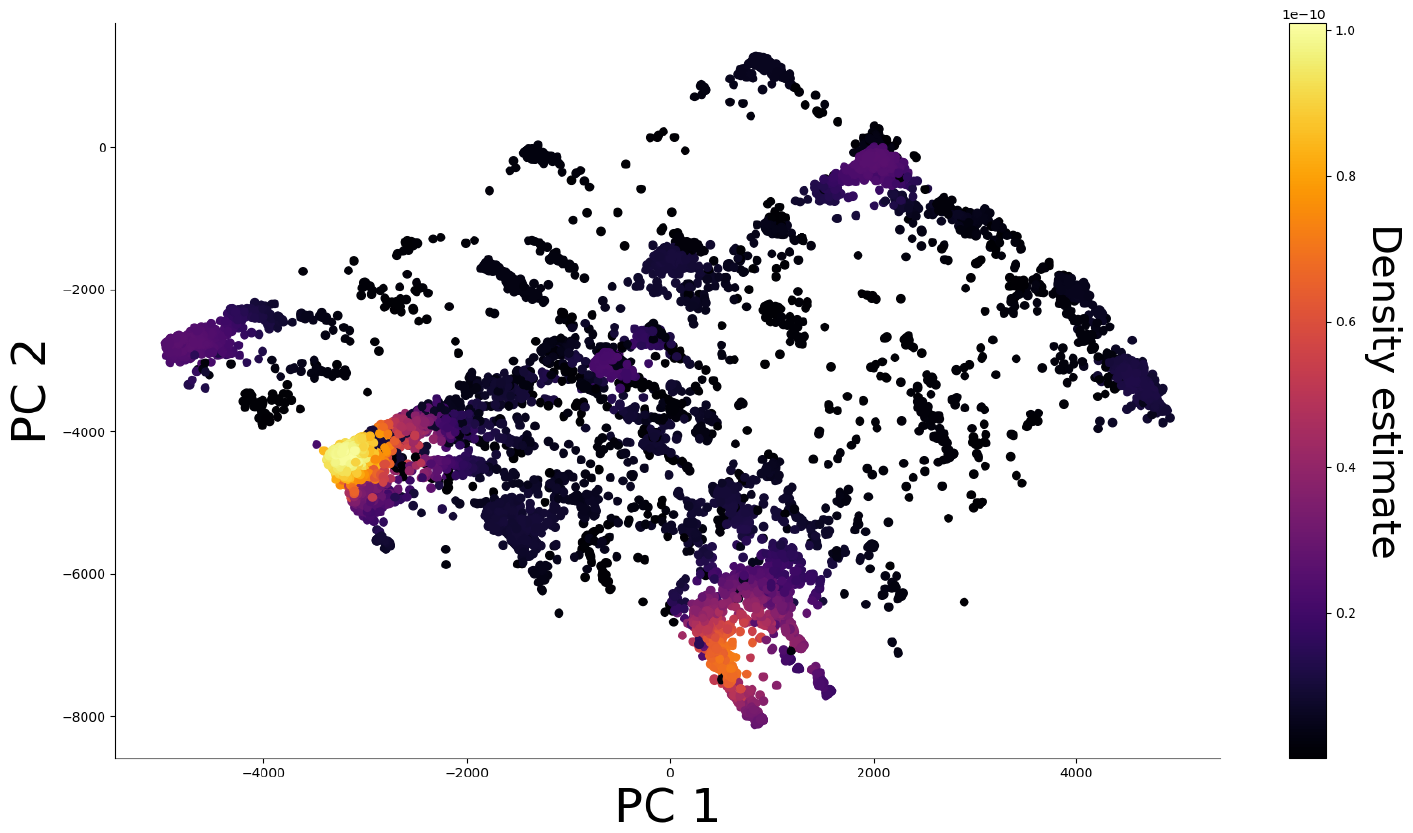
Individual MCMC-samples can be scattered in 2D principal component (PC) space, to find regions of high probability. To make those regions more salient, we colour individual samples according to a Gaussian density estimation (the density estimation occurs in 3D PC space). We can see, that there are multiple modes of varying importance, with one mode being particularly dominant.

In doing this, we found that the posterior for a number of animals wanders itinerantly between different modes, reflecting true uncertainty. These modes are distinct solutions and should not be blended. To isolate them, we performed Gaussian density estimation in the 3D PCA space to identify the ones that were most prevalent, as the regions of highest estimated density. We used this clustering to select samples *j∈ 𝒥* ^*η*^ that were sufficiently similar as to comprise an individual mode 𝒥 ^*η*^. For now, we did this by hand; however, the process could be made more formal by fitting a mixture of Gaussians to the posterior, and then selecting samples around the means of the Gaussians with sufficiently large mixture weights.

Next, we sought to understand how trials within that mode were co-assigned to states. To do this, we averaged the co-occurrences 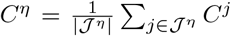, and treated 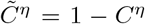 as a distance matrix, where trials were close if they share a state in most samples in the mode. We then performed hierarchical clustering on 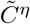, using as a cluster distance 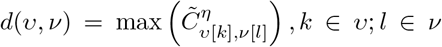, which took as distance between clusters the maximum distance between any two trials in the clusters *u* and *v*. The result of the hierarchical clustering was a tree on the individual trials, and we could extract a clustering by cutting the tree at a certain level. Cutting at, e.g., 0.6 means that we only have clusters in which every trial was explained by the same state in at least 40% (1*−* 0.6) of the samples. For most of our plots we cut at 0.95, which empirically returned good results. Even though this meant that trials needed to use the same state in only 5% of samples to be in one cluster, most trials were assigned to the same state much more often, see **Fig. 11. Fig. 11** also shows a number of alternative clustering from different thresholds, demonstrating that there is little change across a wide range of thresholds: The 95% threshold leads to 8 states with 100% trial coverage, an 80% cutoff leads to 9 states and 99.92% coverage, a 50% cutoff gives 12 states with 98.77% coverage, and lastly a threshold at 20% gives 15 states and 95.27%. We can thus see that low criteria led to trials becoming unassigned and some states splitting apart, which is why we chose a rather high cutoff. A further verfication that the procedure and its threshold gave a faithful representation of the collection of samples comes from comparing the overall solution against individual solutions from single samples. Empirically, these did indeed align.

**Fig. 11.**
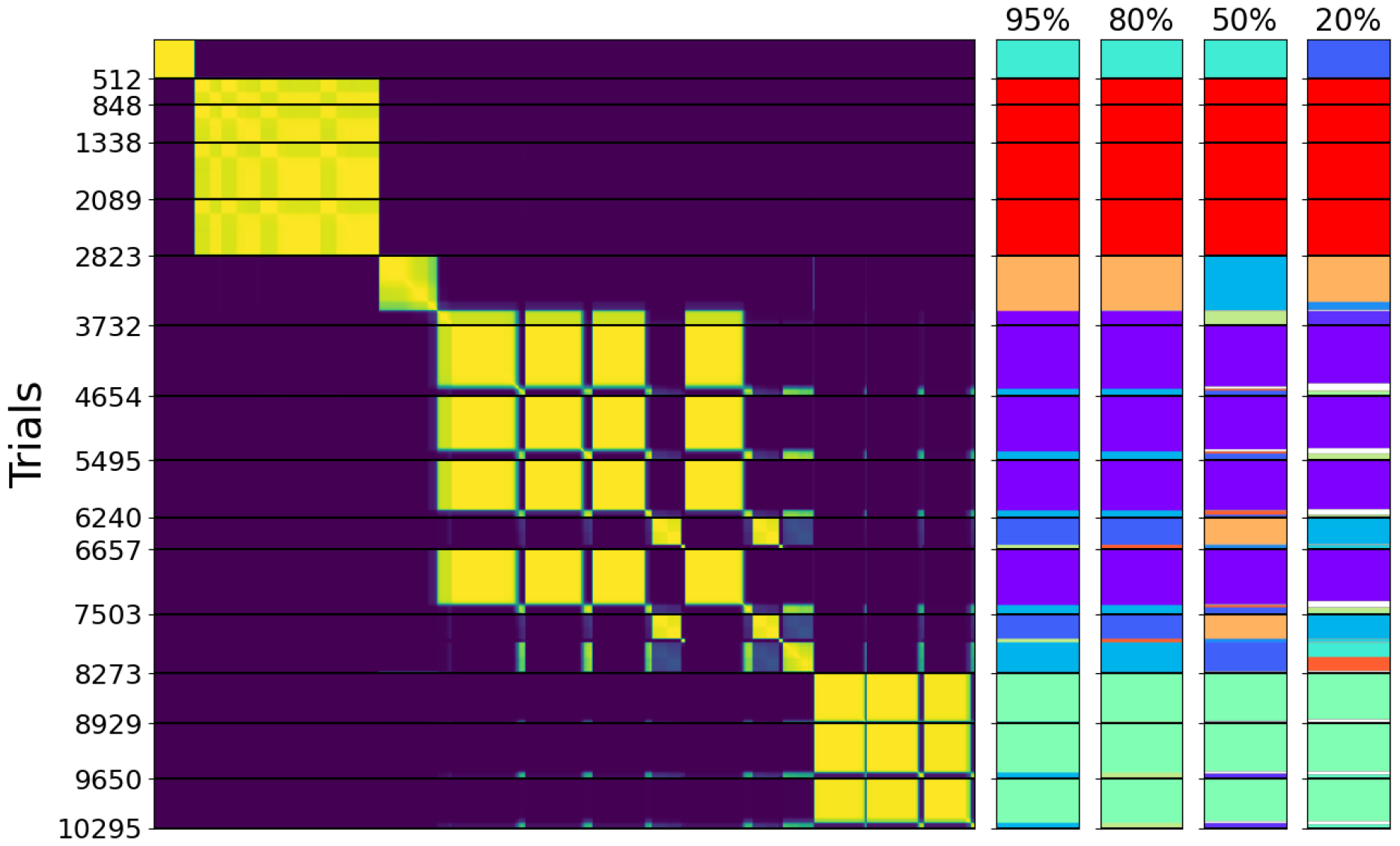
Consistency matrix *C*^*η*^ of the animal seen in **Fig. 2**, with different state assignments, based on cutting the hierarchical clustering tree at different levels, as noted above the colouring assignment. The ticks on the left mark session boundaries, with the numbers indicating the total number of trials so far. State colour on the bars to the right was determined based on the ranking of how many trials are assigned to the state, therefore a colour change need not be a major change in state assignments. As can be seen, state assignments are robust in a large range of cutoff values, with large states staying particularly consistent. Most of the change comes from the splitting of smaller states, and some trials losing a state assignment altogether (a state needed at least 40 trials to be coloured; if the state of a trial had less than that, we colour it white).

The states we show are therefore defined at heart by sets of trials. To compute PMFs, we first considered a single MCMC-sample, and noted which states it assigns to the trials within this set on a session-by-session basis (though each individual trial only had one state assigned in a single sample, for the whole set of trials it usually will not just have been a single state, due to random fluctuations, but mostly a single state). We turned the psychometric weights of these states into PMFs, over which we then averaged (in a weighted manner, considering how often any state occured in the set of trials). For a single sample, this resulted in an average PMF of that state for each session. This then got averaged across samples within a cluster (evenly over all selected samples of a mode), to obtain the ultimate result.

To determine how closely a single trial is connected to its assigned state, we averaged the proportions of samples in which it was in the same state as all the other trials assigned to this state. That is, for a given trial *t*, we took a row of the consistency matrix 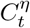, and considered only the entries corresponding to other trials within the state under consideration. We then averaged over those entries, yielding the average proportion of co-assignment. We think of this as a proxy to the posterior over which state a trial is assigned to, as shown in **Fig. 3** (with the caveat that we only determined how strongly a trial is connected to its state, not how much it is connected to other states).

### 6.4 Psychometric type classification

We observed by eye that the psychometric functions (PMFs) that the model found for the behavioural states had a tendency to fall into one of three characteristic classes: flat (type 1), half-tuned (type 2), and fully-tuned (type 3). However, the boundaries between the classes were blurry, so we sought an objective distinction, recognizing its inevitable arbitrariness.

The measure we used in the main paper is the mean reward rate implied by the PMF on easy trials (100% or 50%), ignoring the effects of perseveration (and the debiasing protocol). We chose the reward rate, since this tends to grow as the animals proceed from ignorance to competence. We chose to assess only the easy trials since early PMFs were not defined on the lower contrasts (since these stimuli were not presented). **Fig. 12** shows the distribution of such reward rates across all states. It is apparent that there is a rather clear grouping of PMFs with reward rates below 0.6, defining type 1. The boundary between types 2 and 3 is less evident, implying that edge cases will be hard to assign. The threshold reward rate of 0.78 served reasonably, as evidenced in **Fig. 4**.

**Fig. 12.**
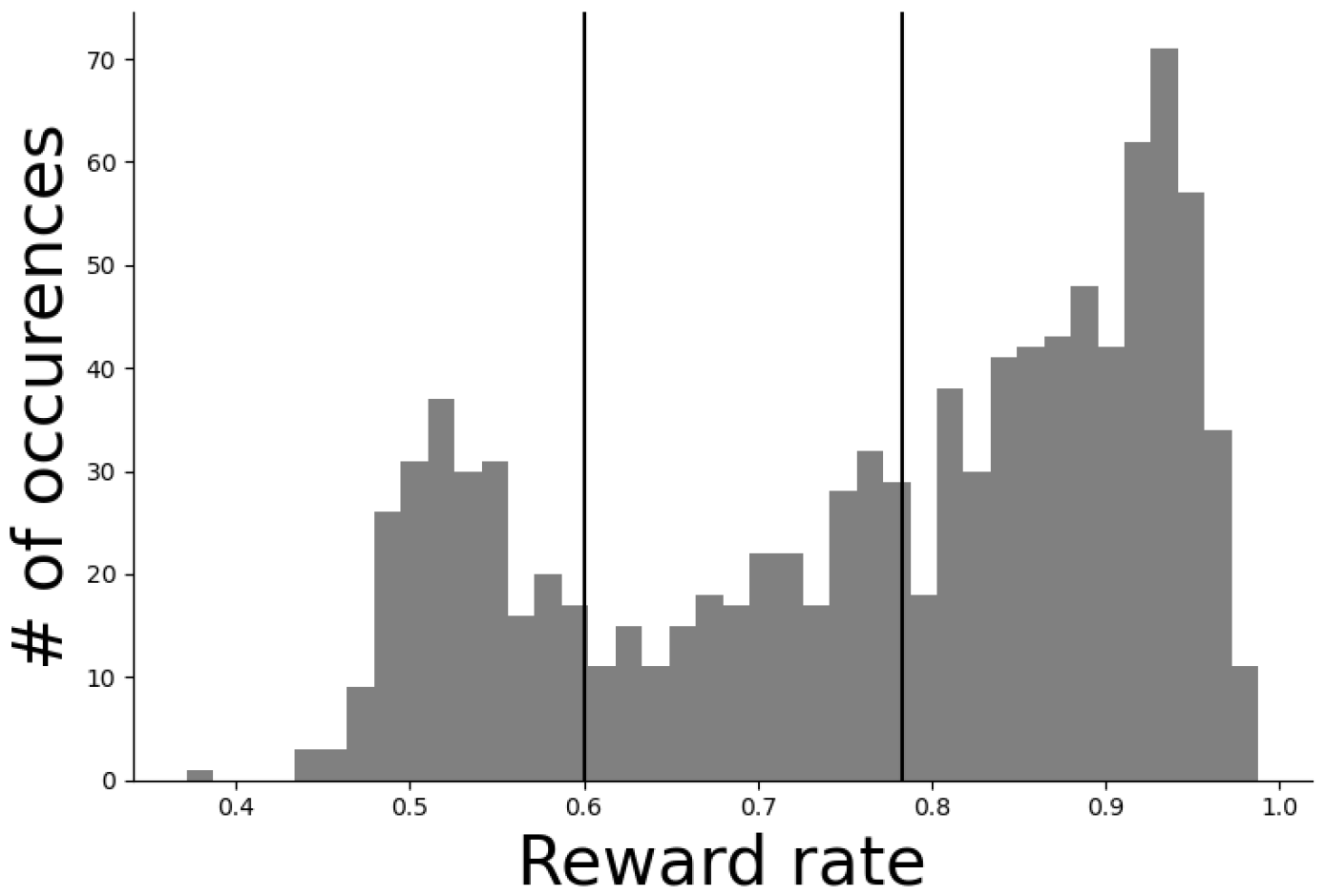
Histogram of the mean reward rates on easy trials of the PMFs of all states, at the moment they first appeared. Vertical lines indicate the boundaries we used to classify states into the three types. The boundaries we drew do align with points of low density in the histogram.

### 6.5 Bias training analysis

As elaborated, the learning steps that were exhibited by the animals seem rather independent, without strong patterns across the stages. This becomes even clearer when considering the next period of learning, the biased training: The basic task stayed the same, but instead of contrasts appearing equiprobably left or right, there were now unsignalled alternations between blocks, lasting 20-100 trials following a truncated exponential distribution, during which a contrast was 80% likely to appear on one side versus 20% on the other. This was of course particularly helpful for 0% contrasts, on which an animal could now reach a much higher reward rate than chance, given a suitable block inference mechanism (a detailed analysis of their actual algorithm was performed in Findling et al. (2023)). To finish this part of training, mice had to show that their behaviour was sufficiently modulated by the current block. Quite surprisingly, the number of sessions it took them to achieve this shows no correlations to any aspect of pre-bias training (Correlation to type 1 duration: Pearson’s r=-0.08, p=0.36; to type 2 duration: Pearson’s r=-0.11, p=0.22; to type 3 duration: Pearson’s r=-0.1, p=0.29; to total pre-bias training: Pearson’s r=-0.13, p=0.15). This suggests that learning about the biased blocks tapped into yet another type of skill, unaddressed by the requirements of the pre-bias protocol.

### 6.6 Model recovery

We tested the model and our inference procedures by fitting to data for which the ground truth was available. For this we instantiated all the random variables of the model to specific values and generated responses from it. This was performed for multiple different variable settings, to assess the accuracy of the fitting procedure in all relevant regimes, and using input data (i.e. contrast sequences) from actual training trajectories. The data generated this way were processed in exactly the same way as those of the IBL mice.

We paid particular attention to assessing the strength of the inductive biases of the inference procedure - particularly in terms of the number of states it inferred (given that this could be potentially unbounded) and the degree of change between sessions (since slow and fast state changes could interact). We tested multiple settings in which all the data were actually generated from a single state, to test whether the model would incorrectly split behaviour into multiple states. In one setting, the psychometric weights of the state stayed constant throughout all sessions, in the other the weights gradually evolved from poor performance to proficiency (at constant steps of a magnitude that corresponds to a variance of 0.0311, the variance of the fitting procedure was still 0.03). Both fits recovered their ground truth successfully, explaining virtually all trials with a single state, as can be seen for the example of the changing state in **Fig. 13**. We also tried a variation of the latter situation, in which the psychometric weights changed in (proportionally smaller) steps on every single trial, rather than all at once at a session boundary (as the model assumes). This, too, was recovered by the model with only one state (which we consider the best possible solution, given that the generative process was outside the model class).

**Fig. 13.**
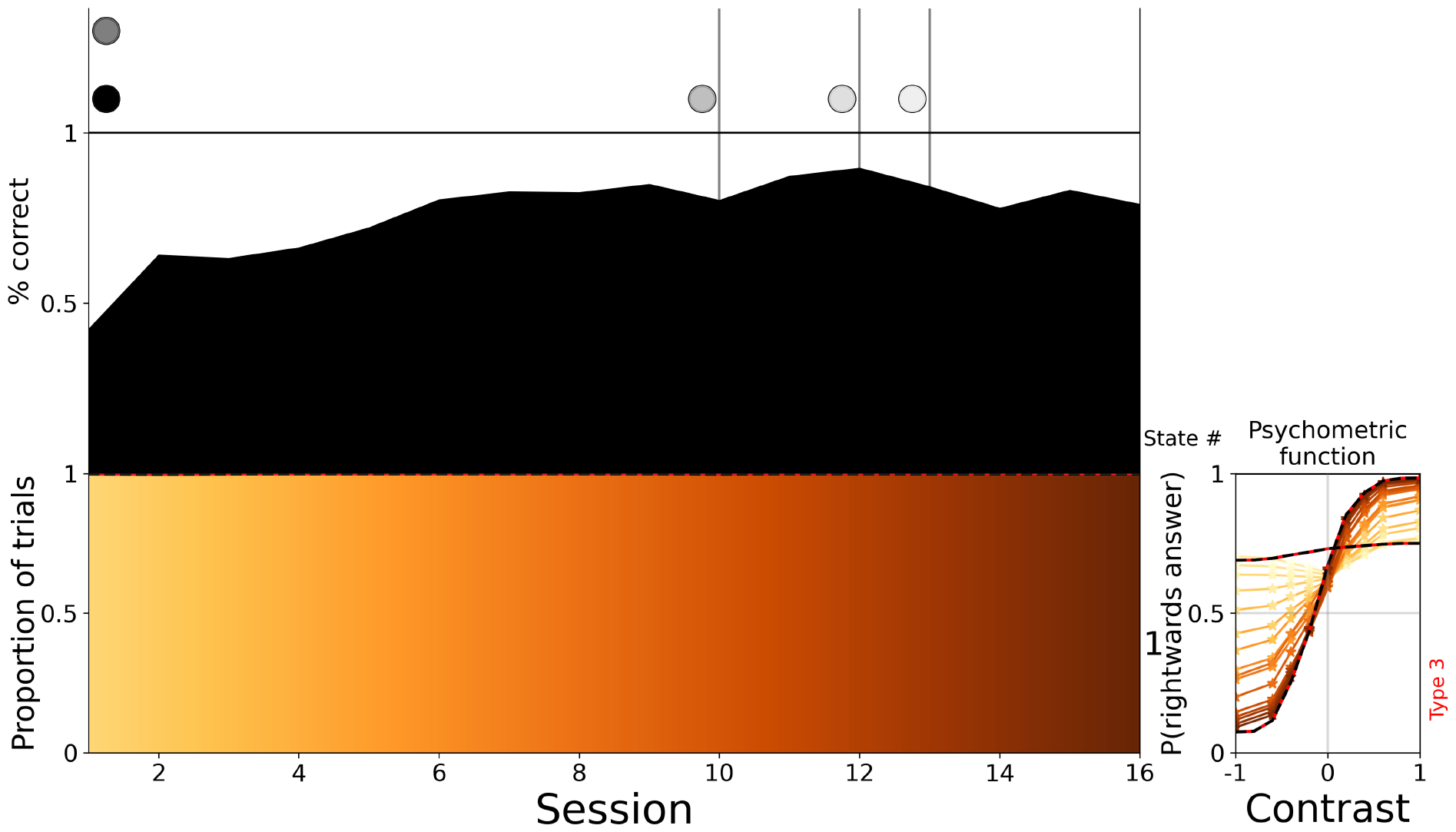
Model recovery where behaviour was generated using only one state, with a changing PMF. The red and black outlines indicate ground truth, which was almost perfectly recovered. For the changing PMF, we only show the first and last ground truth (the initial PMFs are not incorrectly recovered for low contrasts, but are just not learnable due to the limited contrast set during this period)

**Fig. 14.**
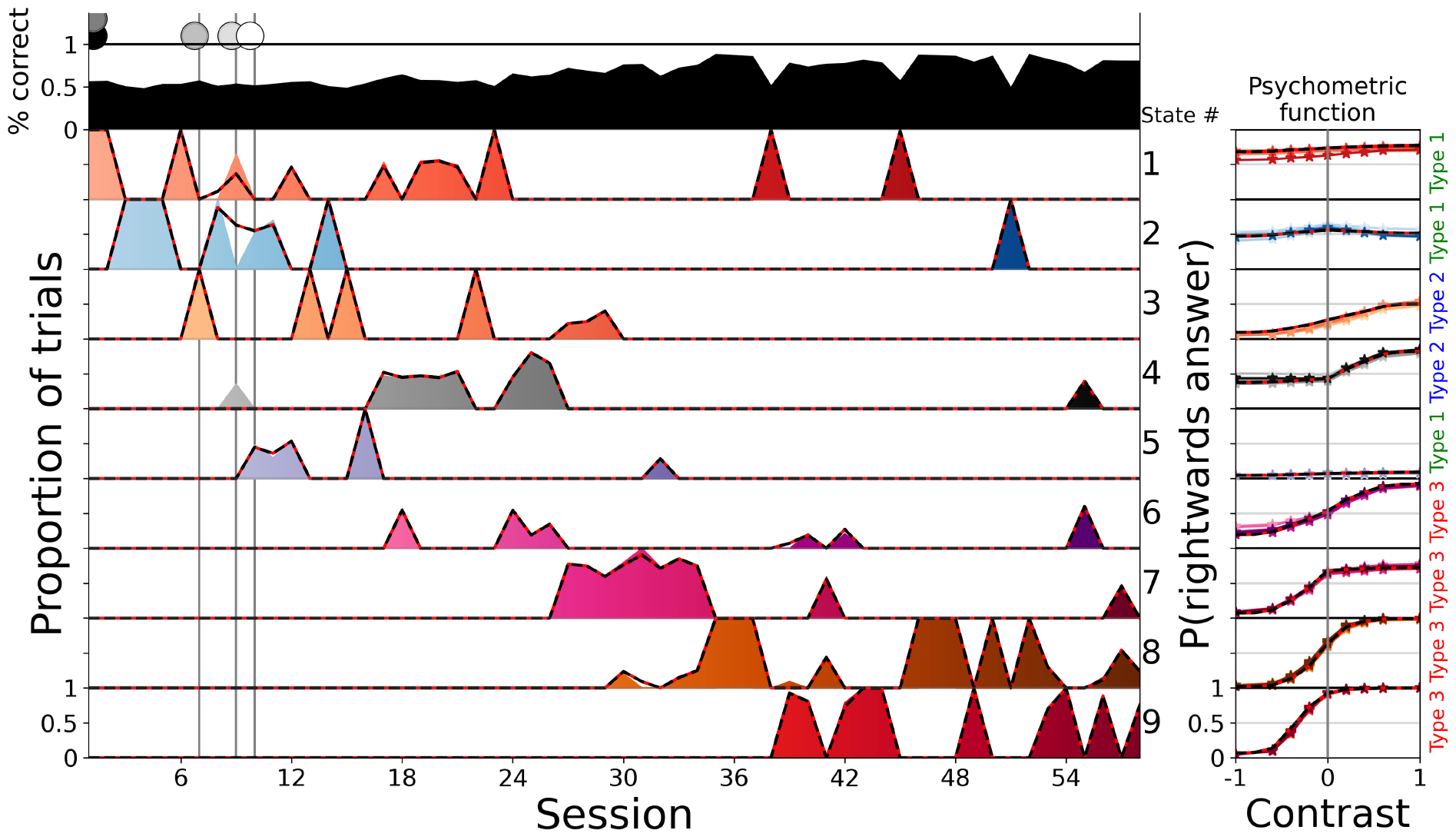
Model recovery where behaviour was generated using 9 different states, on a large number of sessions. The red and black outlines indicate ground truth, which is recovered close to perfectly, in particular correctly recovering the number of states. On session 9, state 2 is incorrectly split between state 1 and 4, the only major flaw.

We also successfully recovered settings from 2 to 9 states, with and without session-to-session variation on the weights, with strongly varying trial proportions between the different states (see **Fig. A18**), and of varying overall training lengths (particularly to test whether long training trajectories lead the model to impose fewer states, making more use of the slow process), as seen in **Fig. 14**. The model was also tested on a setting with completely implausible PMFs, but with the added difficulty of having a larger number of states active within each session, **Fig. A19**. This, too, was captured accurately. These successful recoveries suggest that the model is able to uncover states that truly correlate with distinctly different modes of behaviour in animals.

#### Data access

Please follow these [instructions](https://int-brain-lab.github.io/iblenv/notebooks_external/data_download.html) to download the data used in this article. Use for example the following code snippet to download the data using Python.

~~~
from one.api import ONE
import re
# use password as indicated on the website
one = ONE(base_url=‘https://openalyx.internationalbrainlab.org‘, password=‘*****’)
regexp = re.compile(r’Subjects/\w*/((\w|-)+)/_ibl’)
datasets = one.alyx.rest(‘datasets’, ‘list’, tag=‘2023_Q4_Bruijns_et_al’)
# extract subject names
subjects = [regexp.search(ds[‘file_records’][0][‘relative_path’]).group(1) for ds in datasets]
# reduce to list of unique names
subjects = list(set(subjects))
for subject in subjects:
     trials = one.load_aggregate(‘subjects’, subject, ‘_ibl_subjectTrials.table’)
     training = one.load_aggregate(‘subjects’, subject, ‘_ibl_subjectTraining.table’)
     # save data
~~~

## Acknowledgments

We thank Jonathan Pillow, Anne Churchland, Alexandre Pouget, Charline Tessereau, Zoe Ashwood, Nicholas Roy, Iain Murray, Scott Lindermann and the Behavioural Analysis and Theory working groups of the International Brain Lab for discussions. This work was funded by the Simons Foundation (SAB & PD under ID: 552343, and the IBL in general), the Max Planck Society (SAB, PD), the Wellcome trust (SAB, IBL, PD, ID: 216324), and the Alexander von Humboldt Foundation (PD).

## Authors’ Contributions

SAB and PD initiated the project. SAB and PD developed the modelling framework, which was implemented by SAB. KB, ICL, PYPL, GTM, NJM, JN, AP, NR, KZS, AEU collected the data. SAB and PD performed the data analysis. The paper was jointly written by SAB and PD

## Appendix A Extended Data

**Fig. A15.**
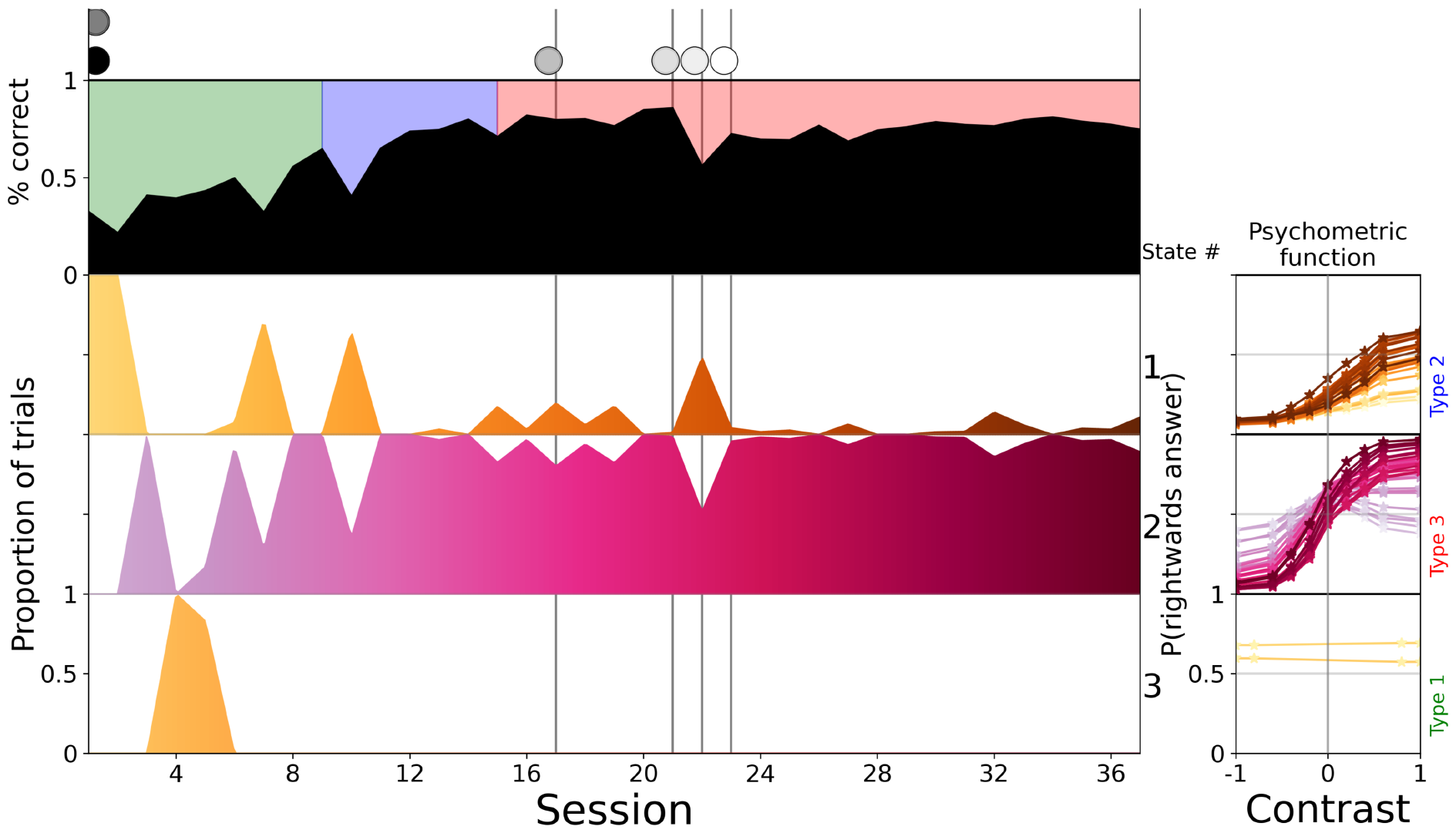
State overview for a mouse with a larger number of sessions. As showcased here, especially for these long training trajectories the model sometimes took some states all the way from uninformed to proficient behaviour, through the slow change process alone.

**Fig. A16.**
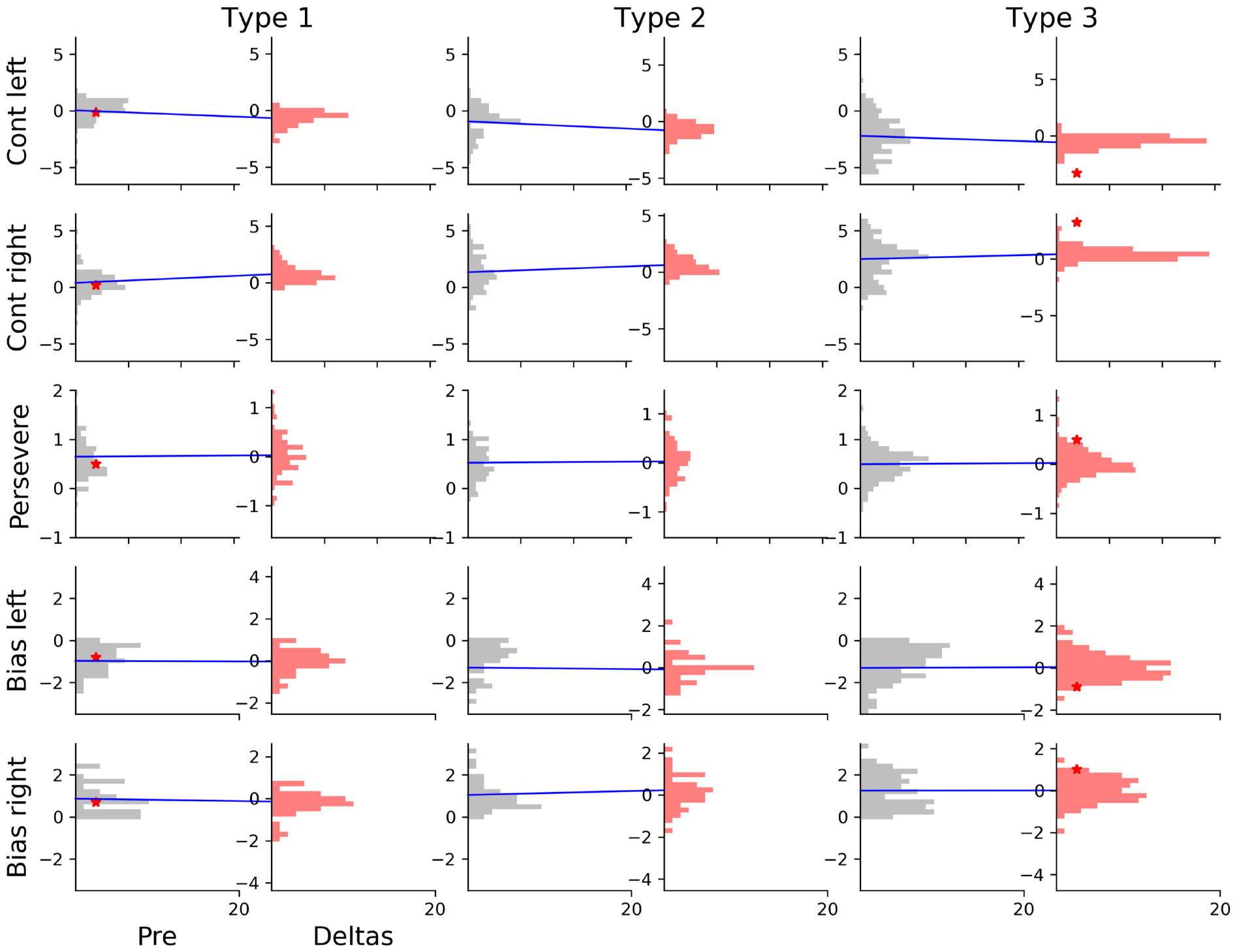
Distributions of initial weights of states and how they changed across their lifespan, split by types. The red distributions to the right show the weight change, not the final distribution, to highlight the change through the slow change process. The blue lines connect the means of two related distributions (we use the mean of the initial distribution as the 0 point of the delta distributions). The bias is split as in **Fig. 6**, and the x-axis is shortened due to this, though the x-ticks are at the same distances across all plots. Red stars mark the average weight of the first and last state of every animal, as in **Fig. 6**

**Fig. A17.**
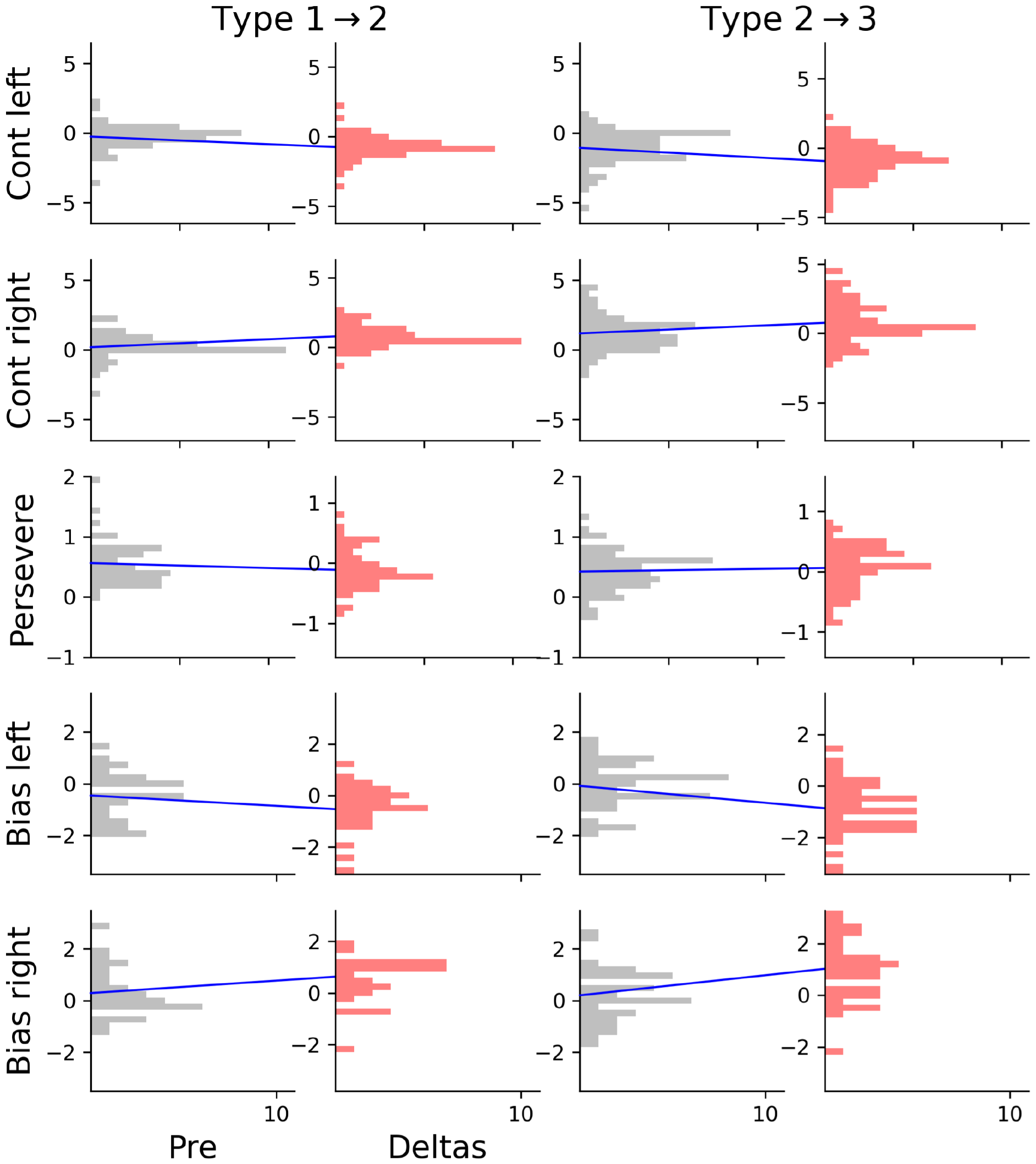
Distributions of weights of the closest previous states and how they differed in the first introduced state of a new type. The red distributions to the right show the weight difference, not the final distribution, to highlight the change through the fast process. The blue lines connect the means of two related distributions (we use the mean of the previous distribution as the 0 point of the delta distributions). The bias is split as in **Fig. 6**, and the x-axis is shortened due to this, though the x-ticks are at the same distances across all plots.

**Fig. A18.**
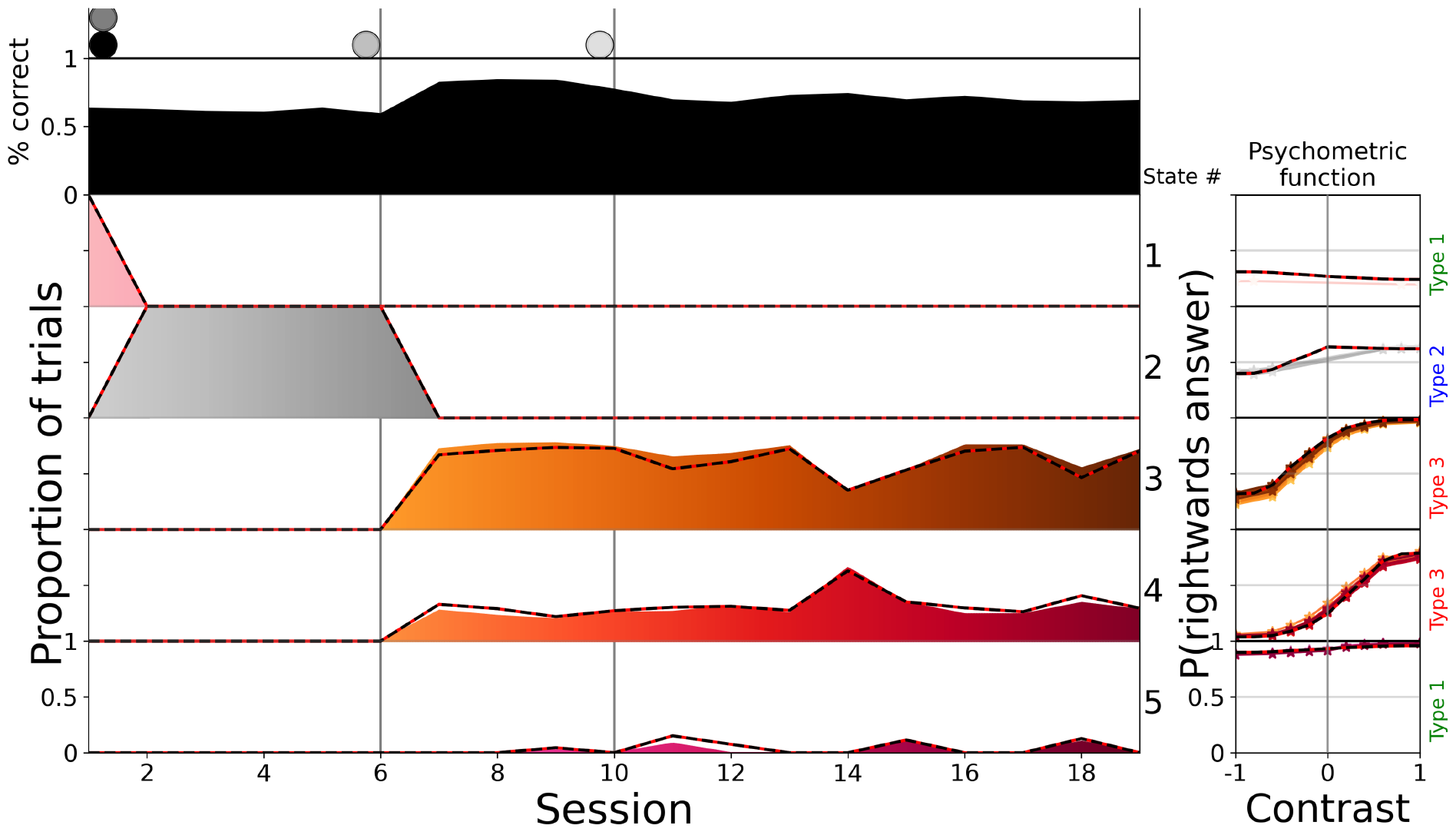
Model recovery of 5 states in a typical progression, the red and black outlines indicate ground truth. This example shows the model’s ability to distinguish between similar states (3 and 4) within a session. A small number of trials got assigned incorrectly between states 3, 4, and 5, in particular the rare state 5 missed some trials.

**Fig. A19.**
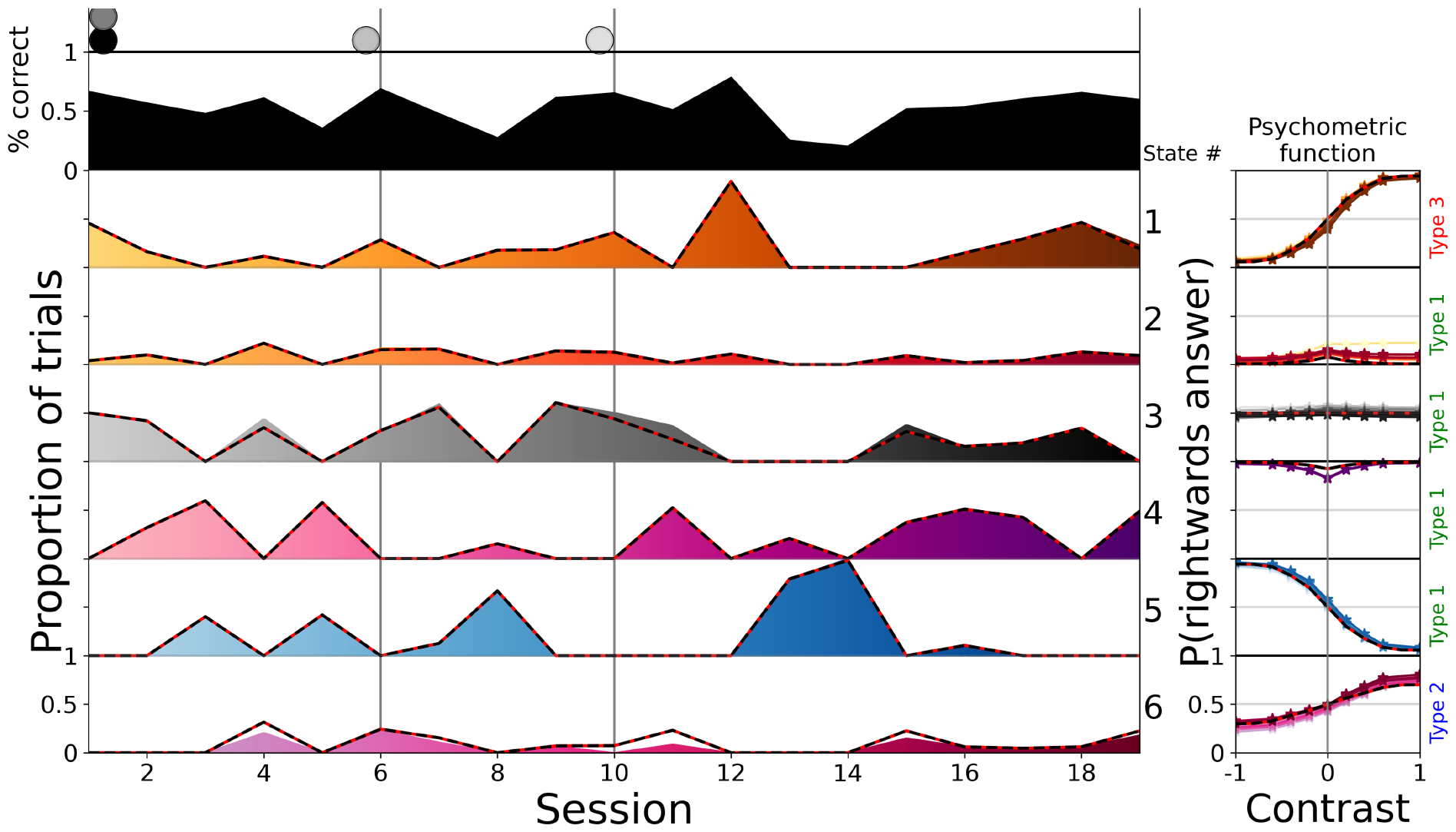
Model recovery of 6 states with some unusual PMFs, which made the states more distinguishable, but the recovery is made more difficult by the fact that many states co-occured in single sessions. The red and black outlines indicate ground truth. The model found the correct number of states for this recovery and almost flawlessly discovered the boundaries between the states in the sessions. Only trials of state 3 and 6, which were the most similar (and state 3 had the highest variance in its responses), were sometimes noticeably misplaced.

